# Neurospheres from primary rodent brain cells to probe the 3D organization and function of synapses

**DOI:** 10.64898/2026.03.19.712855

**Authors:** Benjamin Chauvineau, Adèle Drouet, Charles Ducrot, Léa Bonamy, Tiffany Cloatre, Louis Hurson, Jérôme Baufreton, Jean-Baptiste Sibarita, Olivier Thoumine

## Abstract

To improve our understanding of synapse assembly, there is a need for robust, easy-to-use, and physiologically relevant *in-vitro* models allowing the controllable formation of neuronal contacts in a reasonable time, whose structure and function can be investigated using advanced microscopy. To address this challenge, we engineered 3D cultures from rodent dissociated hippocampal cells, that spontaneously assemble in low attachment U-bottom wells into compact spheroids of reproducible dimensions (100-300 microns), determined by the number of seeded cells. These neurospheres contain a mix of neurons and glial cells and grow over time in culture, through the combination of cell proliferation and neurite extension. Neurospheres were immunostained in fluid phase, and/or sparsely electroporated for the multi-color visualization of synaptic proteins. Neurons extend an elaborate network of axons and dendrites, forming within 2 weeks numerous excitatory and inhibitory synapses identified at the structural level by confocal and electron microscopy, and at the functional level by electrophysiology. Periodic calcium oscillations throughout neurospheres further highlight network activity. Finally, we demonstrate the potential of neurospheres to study synaptogenesis by modulating and visualizing the adhesion protein neuroligin-1. Overall, neurospheres represent a standardized and cost-effective system to study synapse structure and function at high resolution in 3D, that should be quite appealing to the cellular neurobiology community.

## Introduction

The formation of functional synaptic connections between neurons is of utmost importance in brain development, and failure in this process underlies many neurodevelopmental disorders (Bourgeron, 2015; Betancur and Mitchell, 2015). To experimentally investigate synapse organization and function, researchers have engineered over the past decades a wide range of methodologies, including *ex vivo* and *in vitro* culture preparations from wild type or genetically-modified rodents, combined to high-end photon and electron microscopy techniques, chemogenetic and optogenetic modulations, electrophysiological recordings, and functional imaging with fluorescence-based indicators. These advances have led to a major comprehension of the architecture of neuronal circuits and synaptic connectivity in the brain (Kleinfeld et al., 2011; Michalska et al., 2024; Tavakoli et al., 2025). Despite this progress, synaptogenesis remains challenging to study, mainly because of limitations in the biological models used.

The favorite preparation mastered by many laboratories remains primary neurons from rodent hippocampus or cortex, plated on glass coverslips in Banker or mixed configurations (Kaech and Banker, 2006). These 2D cultures allow the polarized growth of axons and dendrites (Dotti et al., 1988), accompanied by formation of synapses in a relatively short time window (typically between days in vitro, DIV10-14) (Cottrell et al., 2000; Chanda et al., 2017). They are well adapted to most super-resolution imaging modalities, except perhaps electron microscopy because of the thinness of the cell layer (Nair et al., 2013; Sun et al., 2018). Moreover, primary neurons can be co-cultured with heterologous cells expressing synaptogenic molecules such as neurexins, neuroligins, SynCAMs, or LRRTMs to decipher the molecular mechanisms underlying pre- and post-synapse maturation (Biederer et al., 2002; Ko et al., 2009; Graf et al., 2004; Scheiffele et al., 2000; Linhoff et al., 2009). However, these simplified 2D cultures remain far from describing complex processes occurring in the living 3D tissue, lack the supporting microenvironment provided by astrocytic processes, and form long-distance connections at unpredictable locations. Micropatterned substrates coated with synaptogenic molecules bring spatial control and high throughput of synapse formation (Czöndör et al., 2013), but they rely on the use of purified proteins that only mimic native receptors in their membrane environment.

On the other hand, flat organotypic slice cultures represent a nice 2.5D system with well-preserved synapse morphology, connectivity, and function (Letellier et al., 2018; Mendez et al., 2010; Stoppini et al., 1991; Toledo et al., 2022; Shipman et al., 2011). However, they are fastidious to produce in large quantities, not easily transfected with plasmids, complex to image in their entirety due to their large spreading surface, display relatively slow synapse dynamics, and are covered with a scar layer hard to penetrate. Human ‘mini-brain’ produced from stem cells is a powerful *in-vitro* system to understand how neuronal circuits develop (Birtele et al., 2024), but at the expense of a high degree of heterogeneity, as well as long and expensive culturing processes. Finally, studying synapse assembly in living rodents is currently out of reach, and faces severe ethical regulations on animal experimentation.

Therefore, to improve our understanding of neuronal connectivity, there is a need for robust, easy-to-use, and physiologically relevant *in-vitro* models allowing for the controllable formation of synapses in a reasonable time, which can be investigated using advanced microscopy techniques. We sought here to combine the easiness of production, genetic manipulation, and labelling of dissociated 2D cultures, to the tissue–like behavior of compliant organotypic slices where neurons are surrounded by supporting glial cells. In this direction, we developed a model of neurospheres formed of primary rodent brain cells, and characterized them extensively using multimodal imaging and functional assays. We first describe the growth and cellular architecture of these 3D objects by immunocytochemistry combined to confocal microscopy. We further reveal the presence of numerous neuronal connections with preserved ultrastructure and function, using electron microscopy, electrophysiology, and live calcium imaging. We then sparsely electroporated neurons with markers of scaffolding and cytoskeletal proteins to highlight excitatory or inhibitory post-synapses as well as dendritic spines, in single neurons. Finally, we used neurospheres for the genetic manipulation and visualization of the cell adhesion molecule neuroligin-1. Overall, neurospheres represent a standardized and cost-effective culture system compatible with existing labelling toolkits and modern imaging techniques, to investigate synapse structure and function at high resolution in 3D. It may be used in the future to screen molecular processes underlying neuronal connectivity in normal and pathological conditions.

## Results

### Size and growth of rat hippocampal neurospheres

We used the traditional cell preparation dissociated from the hippocampi of E18 rat embryos (Kaech and Banker, 2006), but instead of plating cells on polylysine-coated glass coverslips, we pipetted the cell suspension into ultra-low attachment 96-well plates **(Fig. 1A)**. When deprived of the possibility to attach to a substrate, cells grouped together by sedimentation in the U-bottom wells, and formed within 1 day in vitro (DIV) spherical aggregates of highly reproducible morphology and dimension **(Fig. 1B)**. We named these spheroidal objects “neurospheres”. We first measured the diameter of neurospheres (100-300 µm), in relation to the initial number of cells seeded in each well (125, 250, 500 and 1000 cells), and across time in culture (from DIV 1 to 14). As predicted from geometry, the neurosphere diameter (*D_S_*) depends on the cubic root of the number *n* of seeded cells, i.e. *D_S_* = *d_c_ n*^1/3^, where *d_c_* is the diameter of a single cell measured right after dissociation (10.4 ± 1.1 µm, n = 32 cells) **(Fig. 1C)**. For a given number of seeded cells, the neurosphere diameter increased linearly over culture duration, being two times higher at DIV14 than at DIV1 **(Fig. 1D)**, thereby translating into an 8-fold increase in volume. Since hippocampal cells contain a mix of neurons and glial cells, we suspected that this increase in neurosphere size could be partly explained by glial cell proliferation. Indeed, addition of the anti-mitotic agent Ara-C on neurospheres at DIV7 inhibited further growth, although it also seemed to induce some toxicity to the cells. In the rest of the study, neurospheres were formed from 500 or 1000 cells, which were selected for their compactness, lack of aggregation, fast sedimentation, and sufficient size to be detected by eye.

**Figure 1.**
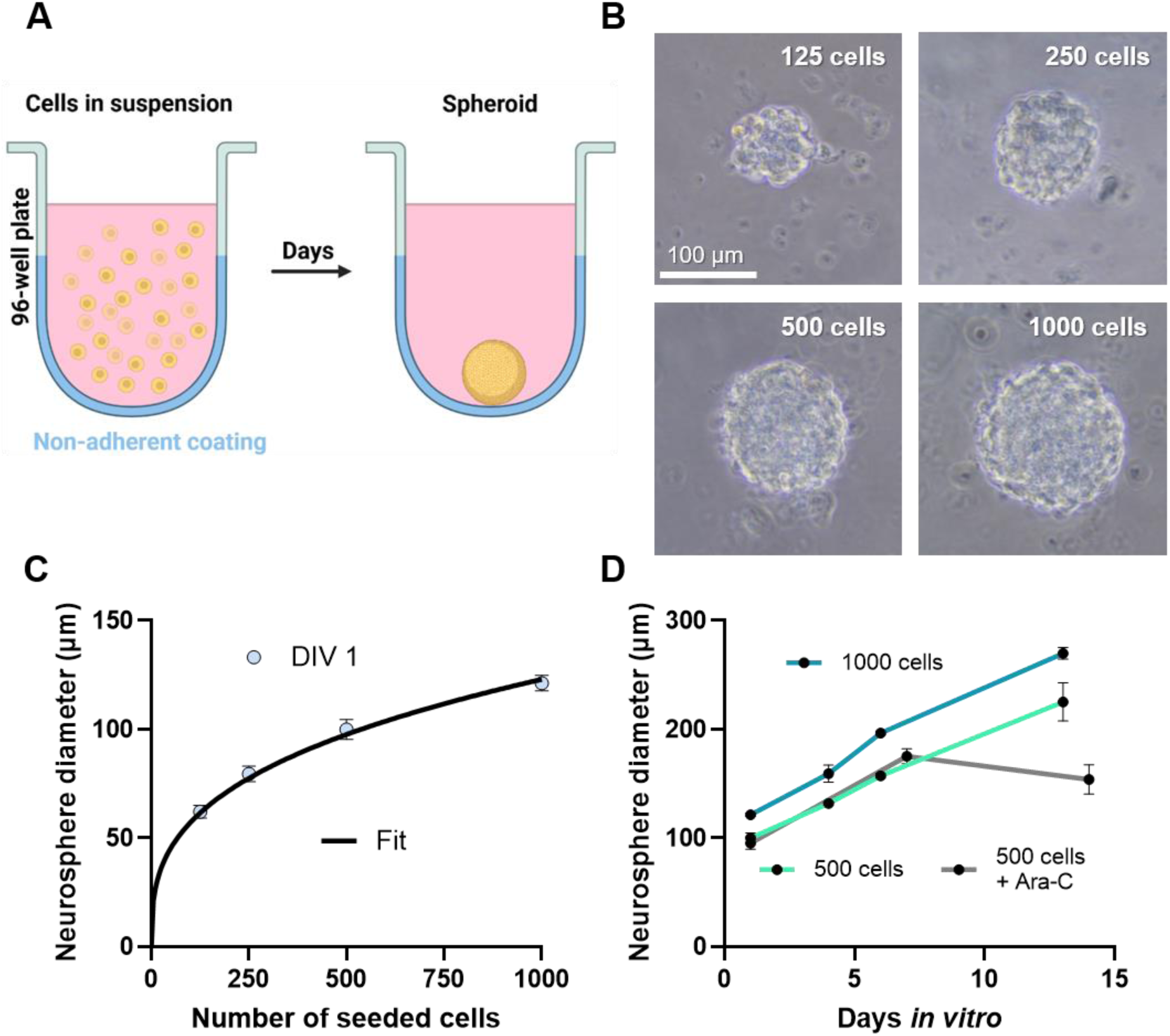
Formation and growth of neurospheres. **(A)** Dissociated hippocampal cells from rat embryos sediment in ultra-low attachment U-bottom wells and form spheroids. **(B)** Brightfield images of neurospheres taken 1 day after being seeded at increasing cell numbers per well. **(C)** Diameter of neurospheres at DIV1 as a function of the initial number of seeded cells per well. The fit through the data (black curve) is a cubit root function. **(D)** Quantification of the neurosphere diameter as a function of time in culture (DIV 1, 4, 6, and 13), for neurospheres initially seeded with 500 cells. In some experiments, Ara-C was added at DIV7, and the neurosphere diameter was measured at DIV14.

### Counting nuclei and distinguishing cell types

To characterize the cells present in neurospheres, we first stained neurospheres initially seeded with 500 cells with DAPI and imaged them in confocal microscopy, at different times in culture **(Fig. 2A)**. We then counted individual nuclei by 3D object segmentation. The number of nuclei per neurophere was slightly below 500 at DIV1 (i.e. some cells migrate on the plastic surface and do not embed in the neurospheres), then increased progressively over time in culture, reaching almost 1400 nuclei at DIV14 **(Fig. 2B)**. Thus, glial cell proliferation does occur, but represents only a fraction of the 8-fold increase in neurosphere volume taking place within two weeks. The second possibility to explain neurosphere growth over culture time is that neurons and glial cells increase their individual volume, e.g. by synthesizing material. Indeed, the apparent cell volume and the distance between neighboring cell nuclei both increased as neurospheres grew with age **(Figs. 2C & S1)**, supporting this hypothesis. To distinguish the cell types populating neurospheres, we immunostained neurospheres in fluid phase **(Fig. S2A)** with the cytosolic neuronal marker NeuN, the astrocytic marker GFAP, or the microtubule associated protein MAP-2, and imaged them in confocal microscopy **(Fig. 2D-F)**. Neurospheres elaborated an extensive network of axons, dendrites, and astrocytic processes, whose overall synthesis likely contributes to the increase in neurosphere volume. The proportion of NeuN-positive cells (i.e. neurons) versus the total number of nuclei quantified in selected confocal planes **(Fig. 2G)**, was around 40% at DIV14. Interestingly, neuronal nuclei were larger and less intense than those of NeuN-negative cells **(Fig. 2H,I)**. These properties suggest that glial cells, contrary to neurons, are in a proliferative state. Indeed, some NeuN-negative cells were clearly identified in the process of dividing. Together, these findings indicate that neurospheres preserve a diversity of cell types which exists in the original hippocampal tissue. Furthermore, neurosphere growth through glial cell proliferation and neurite synthesis is taken as an indicator of the health and integrity of the sample.

**Figure 2.**
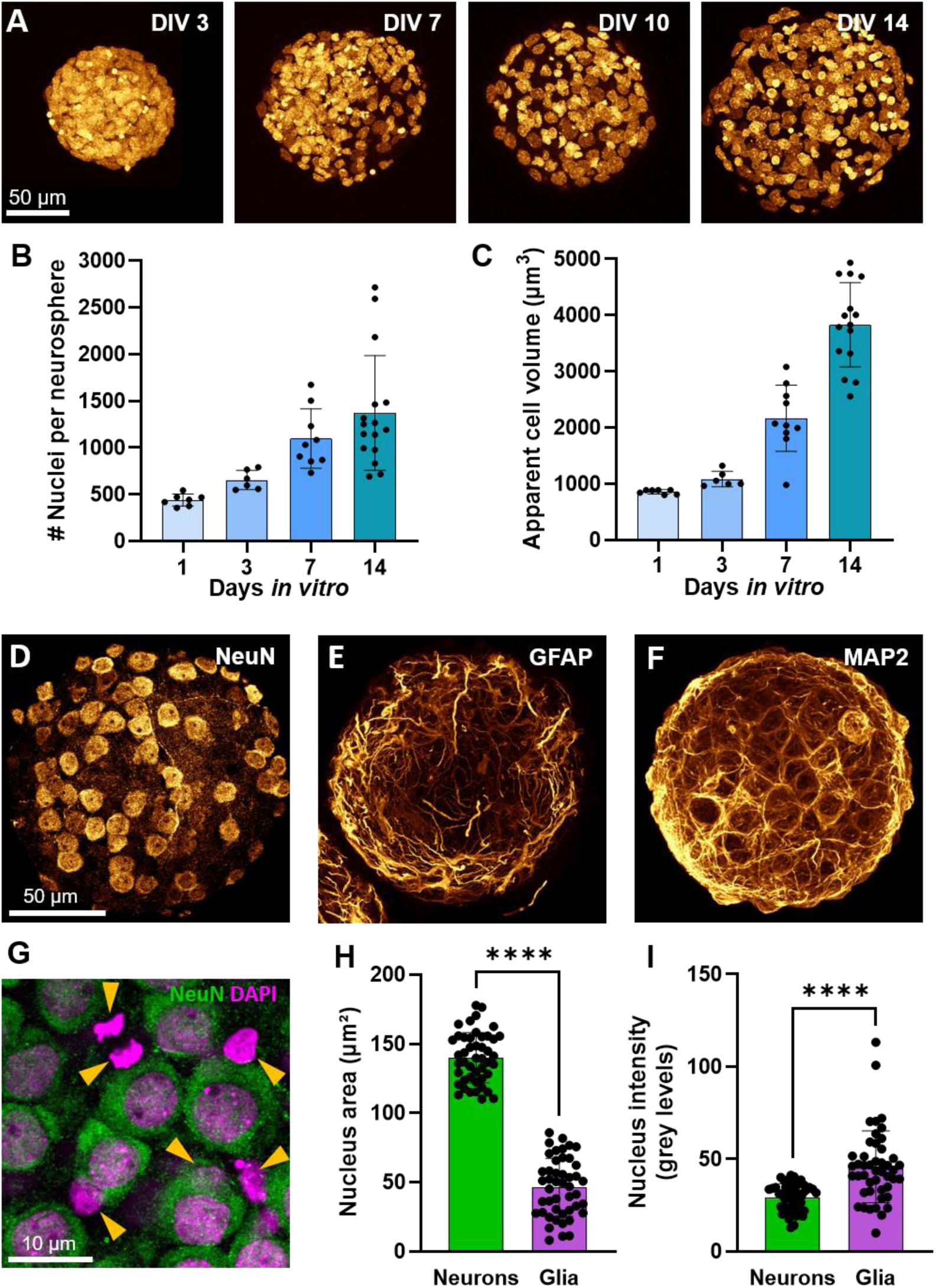
Imaging nuclei and cell types across culture time. **(A)** Maximal intensity projections of confocal image stacks of DAPI staining in DIV 1, 3, 7, and 14 neurospheres initially formed from 500 cells. **(B, C)**. Number of individual nuclei and apparent cell volume, as a function of neurosphere culture time. **(D-F)** Maximum intensity projection images of DIV14 neurospheres stained with antibodies to NeuN, GFAP, or MAP-2, respectively. **(G)** Merged confocal image of NeuN (green) and corresponding DAPI (magenta) staining, at higher magnification. Note the presence of NeuN-negative nuclei of small size, some of them revealing recent cell division (arrowheads). **(H, I)** 2D projected area and relative DAPI intensity of NeuN-positive versus NeuN negative nuclei, respectively (dots in the graph represent individual nuclei). Data are presented as mean ± SEM and were compared by a non-parametric Mann-Whitney test (n = 48 and 46 nuclei per condition).

### Formation of synaptic connections

To assess whether neurons formed *bona fide* synapses between themselves, we immunostained neuropheres with the pre-synaptic markers Synapsin1, VGluT1, or VGAT, together with MAP-2, and imaged them in confocal microscopy. There were many puncta detected along dendrites, indicating the presence of both excitatory and inhibitory synapses **(Fig. 3A,B)**. Using 3D image analysis, we counted on average 579 ± 48 VGluT1 positive puncta and 333 ± 35 VGAT positive puncta per neuron at DIV14 (n = 22 and 8 neurospheres, respectively) **(Fig. 3C)**. The synapsin1 immunostaining increased over developmental age, with a sharp transition between DIV7 and DIV10 **(Fig. 3D,E)**, a period of active synaptogenesis in those cultures (Cottrell et al., 2000; Chanda et al., 2017; Czöndör et al., 2012). Second, we fixed and stained neurospheres with osmium, embedded them in resin, and cut them into ultrathin sections before observation under transmission electron microscopy (TEM) **(Fig. S2B,C)**. We found both asymmetric synapses with visible post-synaptic density (PSD), likely corresponding to excitatory inputs on dendritic spines, and symmetric synapses without detectable PSD, likely corresponding to inhibitory synapses **(Fig. 3F,G)**.

**Figure 3.**
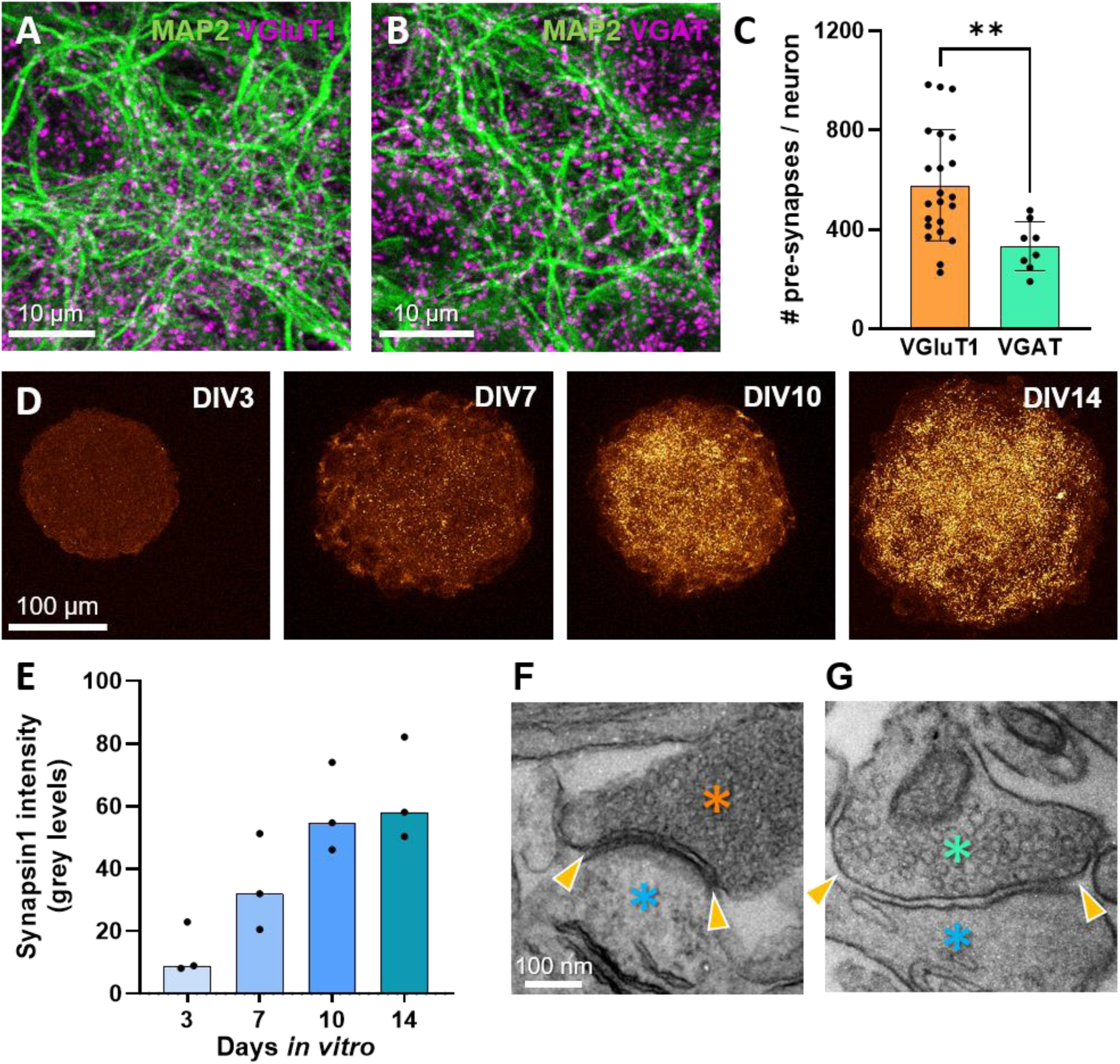
Imaging synapses by confocal microscopy and TEM (A, **B)** Confocal images of VGluT1 and VGAT puncta (magenta), respectively, along microtubules labelled with MAP-2 (green). **(C)** Quantification of the total number of excitatory and inhibitory presynapses in DIV14 neurospheres, normalized by the estimated number of neurons. Data represent the mean ± SEM of 22 and 8 neurospheres, respectively, and were compared by non-parametric Mann-Whitney test. **(D)** Maximum intensity projections of confocal images of neurospheres at DIV 3, 7, 10, and 14, immunostained for synapsin1. **(E)** Graph showing the average Synapsin1 immunofluorescence signal quantified on maximal intensity projection images of neurospheres at varying developmental age. **(F, G)** Representative TEM images of an excitatory and an inhibitory synapse, respectively. Pre-synaptic elements are highlighted with orange and green stars, post-synaptic elements in blue, and the synaptic cleft with arrowheads. Note the presence of a PSD in the excitatory synapse.

### Neuronal excitability and synaptic transmission

To investigate the properties of neuronal excitability and synaptic transmission at the single-cell level, we used electrophysiology. Individual cells in neurospheres were subjected to patch-clamp **(Fig. 4A)**, and their capacity to elicit action potentials and to receive synaptic inputs were measured. Patched cells had electrophysiological parameters in the range expected for neurons **(Table 1)**. In current clamp-mode, increasing depolarization caused neurons to fire **(Fig. 4B)**, showing a positive relationship between the injected current and the frequency of action potentials with saturation at the highest currents **(Fig. 4C)**. In voltage-clamp mode, we first recorded spontaneous EPSCs at -60 mV for 5 min, then spontaneous IPSCs at + 10 mV from the same neurons (Levinson et al., 2005; Letellier et al., 2018), revealing the formation of functional excitatory and inhibitory synapses **(Fig. 4D-G)**. We verified that sEPSCs were mediated by AMPA receptors by adding the drug DNQX, and that sIPSCs were mediated by GABA-A receptors by adding gabazine. The amplitude of sEPSCs and sIPSCs was 39 ± 6 pA versus 93 ± 25 pA (n = 11 and 8 patched neurons, respectively) **(Fig. 4H)** and their frequency was 7.2 ± 1.4 Hz versus 5.9 ± 1.8 Hz **(Fig. 4I)**. These values are comparable to those previously measured in hippocampal dissociated neurons or organotypic slices (Letellier et al., 2018; Haas et al., 2018; Letellier et al., 2020; Kwon et al., 2012), indicating that pre-synaptic and post-synaptic elements are assembled as properly in neurospheres as in the two other culture models. The proportion of neurons that exhibited spontaneous EPSCs and IPSCs was higher when neurospheres were cultured in BrainPhys medium compared to Neurobasal medium **(Fig. S3A-E)**, as previously experienced (Dubes et al., 2022; Bardy et al., 2015). In some neurons, bursts of excitatory synaptic activity lasting around 1-2 sec were detected **(Fig. S3F,G)**. Finally, spontaneous sEPSCs were blocked by addition of the sodium channel blocker tetrodotoxin (TTX), suggesting that they are triggered by action potential-driven glutamate release at pre-synapses **(Fig. S3H-J)**. Therefore, in neurospheres, individual neurons form functional excitatory and inhibitory synapses with typical electrophysiological properties.

**Figure 4.**
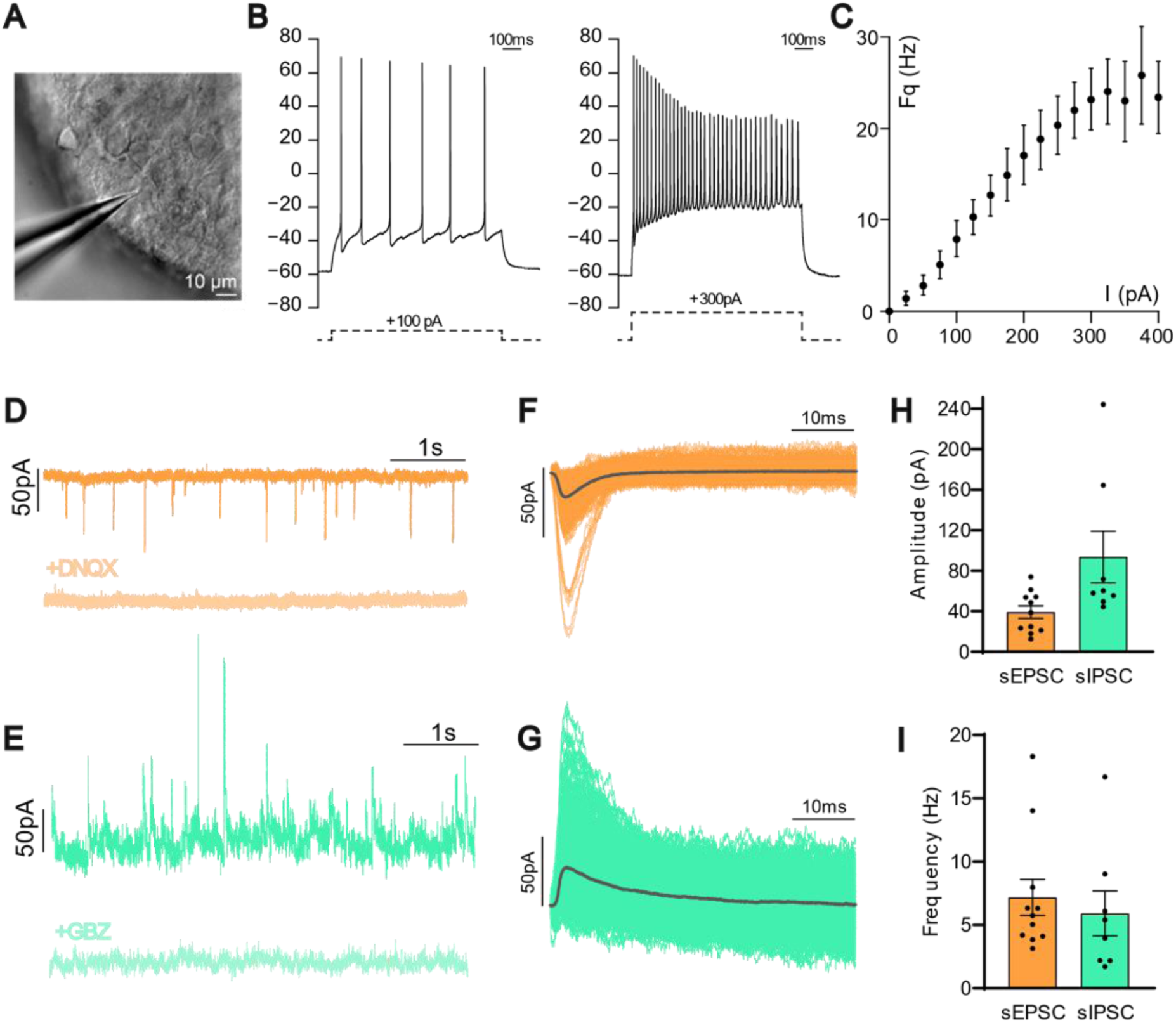
Electrophysiological characterization of neurons present in neurospheres. **(A)** DIC image showing part of a neurosphere with the patch pipette used for electrophysiological recordings. **(B)** Representative recordings of the membrane potential of a single neuron held in current clamp, upon current injection of +100 and +300 pA, respectively. Note the increase in the number of evoked action potentials with the amount of injected current. **(C)** Relationship between the injected current (I) and the frequency (Fq) of action potentials generated by the neurons (mean ± SEM of n = 21 neurons). **(D, E)** Representative traces depicting spontaneous EPSCs and IPSCs, recorded at holding potentials of -60 mV and +10 mV, respectively. In the lower traces, addition of the AMPA or GABA-A receptor antagonists DNQX (20 µM) and gabazine (10 µM), respectively, block the corresponding currents. **(F, G)** Superimposed individual traces of spontaneous EPSCs (orange, 480 events) and IPSCs (green, 932 events), respectively, together with the corresponding average traces (black). **(H, I)** Amplitude and frequency of spontaneous EPSCs and IPSCs, respectively (n = 11 neurons from 9 neurospheres). Data are presented as the mean ± SEM. Dots in population graphs represent individual neurons.

**Table 1:** Electrophysiological parameters.

| Parameter | Mean value | SEM | # cells |
| --- | --- | --- | --- |
| **Excitability** |  |  |  |
| Resting membrane potential (mV) | -61,1 | 1,8 | 20 |
| Capacitance (pF) | 81,5 | 6,7 | 20 |
| Resistance (MΩ) | 262,3 | 25,8 | 19 |
| **Spontaneous synaptic transmission** |  |  |  |
| Rise time EPSC (ms) | 1,4 | 0,1 | 11 |
| Tau EPSC (ms) | 4,3 | 0,4 | 11 |
| Rise Time IPSC (ms) | 2,8 | 0,3 | 8 |
| Tau IPSC (ms) | 15,7 | 1,3 | 8 |

### Spontaneous calcium transients in neurospheres

To assess inter-neuron connectivity at the level of entire neurospheres, we used calcium imaging. Neurospheres previously cultured in BrainPhys medium were loaded with the cell permeant calcium indicator Fluo-4 AM, and imaged live under epifluorescence microscopy. Within a 1-2 min observation period, 82% of DIV14 neurospheres (108 out of 132) displayed spontaneous calcium transients **(Fig. 5A,B)**, while the rest remained silent. The Fluo-4 signals from individual cells at different locations were highly synchronized, revealing a fast propagation of calcium oscillations throughout the whole neurospheres **(Movie 1)**. The peak amplitude of calcium spikes varied between 10 and 150% of the baseline, while the average temporal width was 4.4 ± 0.2 sec (n = 95 pulses analyzed). The frequency of calcium transients ranged between 0.5 and 35 oscillations per min, with two main populations of neurospheres: one displaying high amplitudes and low frequency, the other having low amplitudes and high frequency **(Fig. 5B,C)**. Overall, there was a decreasing relationship between the amplitude and frequency of calcium transients **(Fig. 5D)**. Calcium oscillations were completely suppressed by treatment with TTX **(Fig. 5E, Fig. S4A, and Movie 2)**, demonstrating that they depend on action potential propagation along axons. On the other hand, treatment with 100 µM glutamate to stimulate synaptic transmission, or with 40 mM KCl to induce membrane depolarization, both induced massive increases in Fluo-4 signals among all neurospheres in the field of view **(Fig. 5E, Fig. S4A-D, and Movies 3 and 4)**. These results suggest that calcium transients are mediated by glutamatergic synaptic transmission, potentially depolarizing the membrane sufficiently to open voltage-gated calcium channels (Heine et al., 2008). Finally, we performed calcium imaging of neurospheres at various developmental ages (DIV3, 7, 10, and 14). The measured calcium influx sharply increased between DIV7 and 10 **(Fig. 5F)**, correlating with the synaptogenesis onset. Together, these observations demonstrate fast, synapse-dependent, and short-range functional connectivity throughout the neuron network.

**Figure 5.**
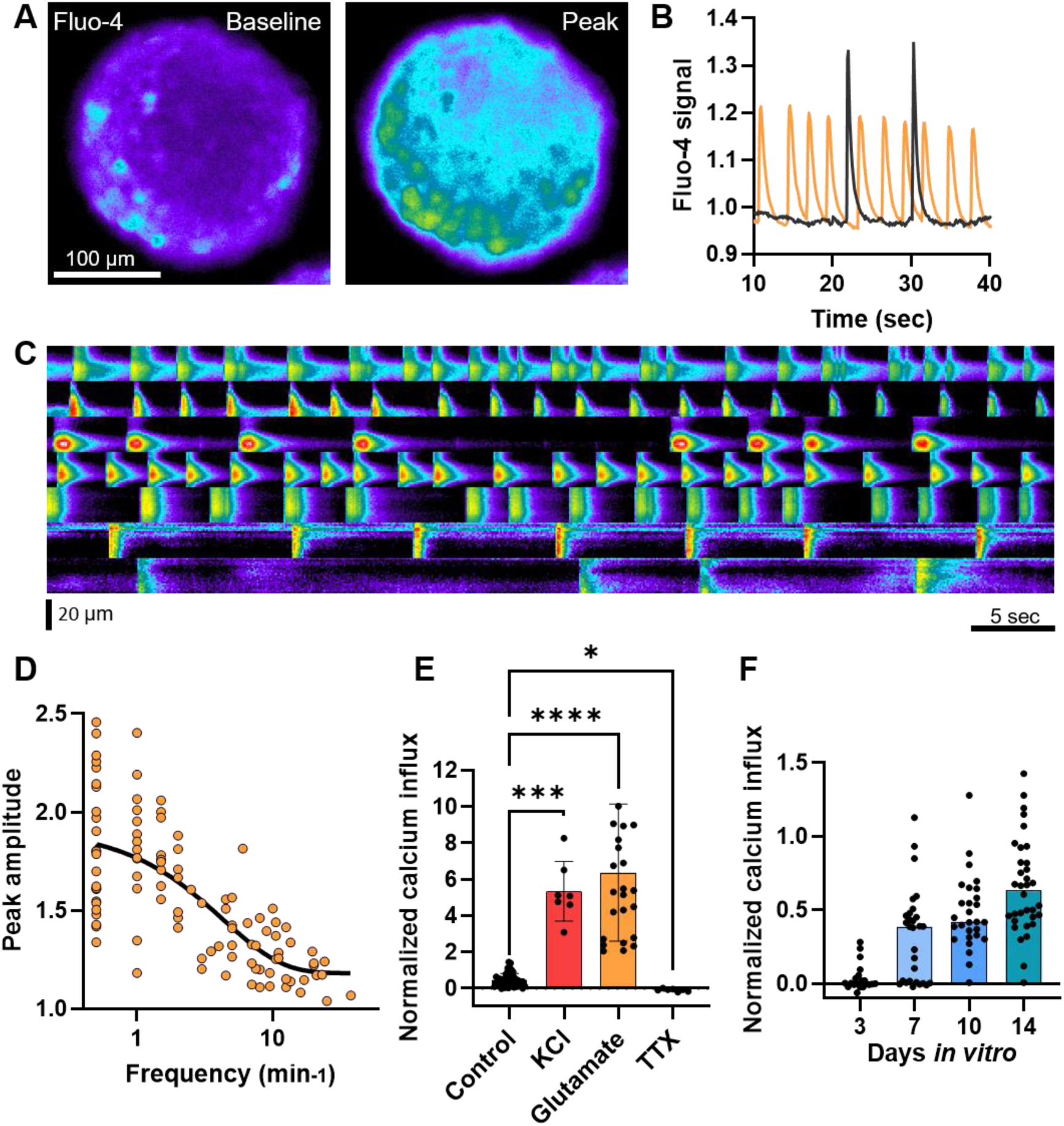
Calcium imaging in neurospheres. **(A)** Epifluorescence images of an entire neurosphere loaded with the cell-permeant calcium indicator Fluo-4 AM, taken at baseline and during a calcium spike (peak). **(B)** Spontaneous calcium transients elicited by two different neurospheres during a 40 sec time period, normalized to the baseline. **(C)** Kimographs of Fluo-4 signals over time for seven independent neurospheres. The baseline was set at 0 grey levels, and the Fluo-4 intensity represents the normalized peak value multiplied by 1000 grey levels. **(D)** Relationship between the frequency of the calcium transients and their peak amplitude, each dot representing a single neurosphere. **(E)** Normalized calcium influx in control condition, or upon treatment with various drugs (KCl, glutamate, or TTX). **(F)** Normalized calcium influx in untreated neurospheres measured at different times in culture (DIV 3, 7, 10, and 14).

### Imaging individual post-synapses at high resolution

To increase our imaging resolution of individual synapses, we used existing electroporation and labeling toolkits for synaptic proteins (Chamma et al., 2016; Chazeau et al., 2015; Fischer et al., 2000). Hippocampal cells in suspension were electroporated right before being seeded into the wells and mixed with non-electroporated cells in order to isolate a sparse population of labeled neurons **(Fig. 6A)**. Neurospheres were then imaged in 3D using confocal microscopy, with or without immunocytochemical amplification of the fluorescent proteins (GFP, RFP…) fused to the proteins of interest **(Fig. 6B)**. Electroporated neurons were preferentially found at the periphery of neurospheres and extended long axons that invaded the whole neurosphere. Neurons electroporated with an intrabody to PSD-95 (Xph20-GFP) (Rimbault et al., 2024) exhibited discrete puncta around dendrites corresponding to excitatory post-synapses, while neurons electroporated with an intrabody to gephyrin (GPHN.FingR-GFP) (Gross et al., 2013) displayed puncta around cell bodies and in dendritic shafts, corresponding to inhibitory post-synapses **(Fig. 6C)**. We quantified the numbers of PSD-95 and gephyrin-positive puncta per electroporated neuron, and found more excitatory post-synapses than inhibitory ones **(Fig. 6D)**, confirming the data obtained for the pre-synaptic markers VGluT1 and VGAT at the population level. By co-electroporating neurons with GPHN.FingR-GFP and Xph20-mRuby, we could even detect both excitatory and inhibitory post-synapses in the same neurons **(Fig. 6E,F)**. Neurons expressing GFP-tagged actin displayed numerous well-formed spines in proximal dendrites **(Fig. 6G)**, with a linear density of 0.82 ± 0.10 spines per µm dendrite length (n = 16 neurons), in agreement with measurements made on CA1 pyramidal neurons in organotypic hippocampal slices or on dissociated cortical neurons (Letellier et al., 2018; Shipman et al., 2011; Kwon et al., 2012).

**Figure 6.**
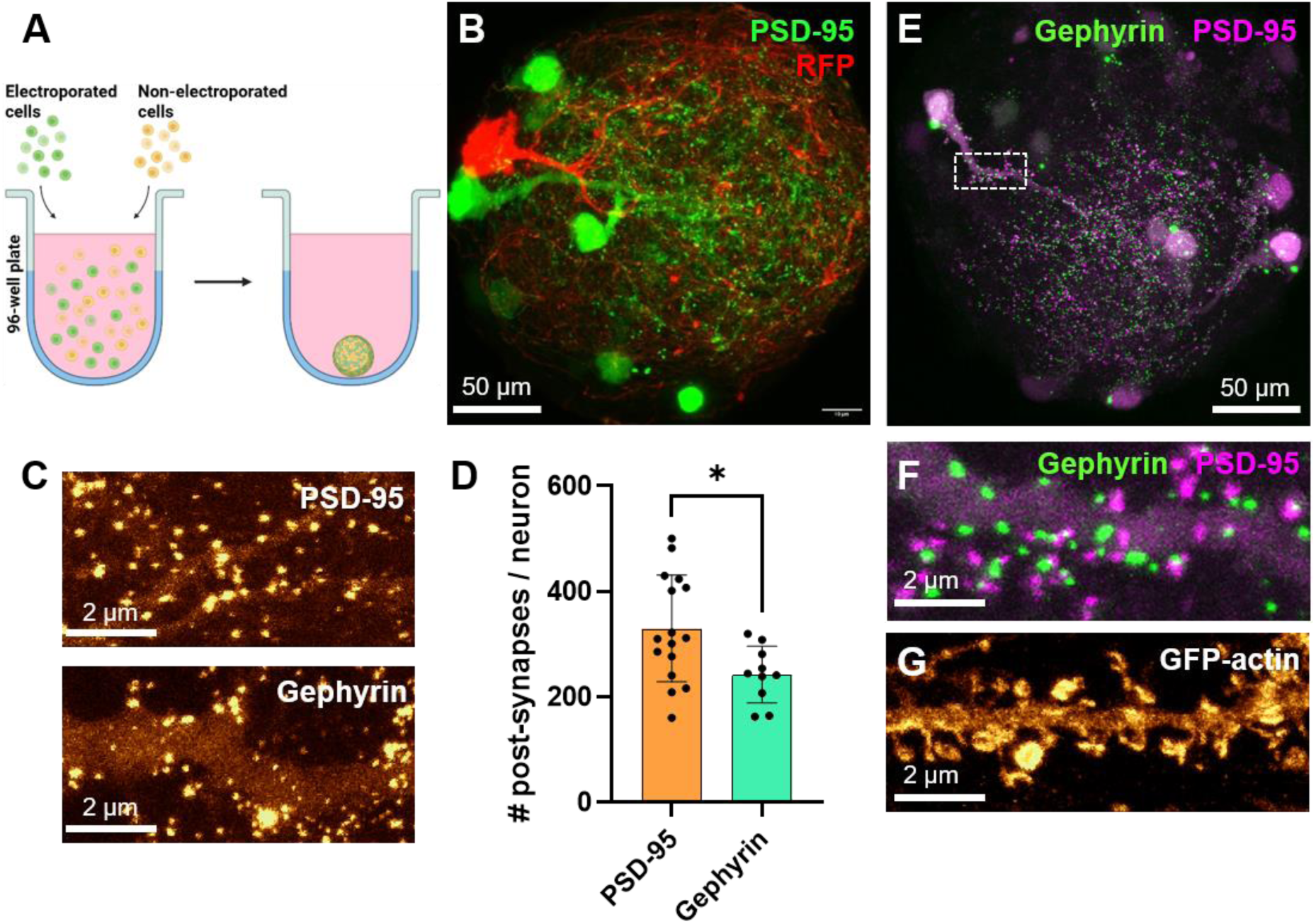
Imaging of post-synapses in sparsely electroporated neurons. **(A)** Neurospheres were formed by mixing non-electroporated and electroporated cells before being plated in U-bottom ULA wells. **(B)** Maximum intensity projection image of a confocal stack of a DIV14 neurosphere in which neurons were separately electroporated with RFP (red) and Xph20-GFP (green). **(C)** Zoomed images on primary dendrites from neurons expressing intrabodies to PSD-95 (Xph20-GFP), or gephyrin (GPHN.FingR-GFP), revealing excitatory or inhibitory post-synapses, respectively. **(D)** Numbers of PSD-95 and gephyrin-positive puncta per electroporated neuron. Data represent the mean ± SEM of 14 and 10 neurospheres, respectively, and were compared by non-parametric Mann-Whitney test. Dots show individual neurospheres. **(E)** Maximum intensity projection of a confocal stack of a DIV14 neurosphere in which neurons were co-electroporated with GPHN.FingR-GFP (green) and Xph20-mRuby2 (magenta), allowing the detection of both excitatory and inhibitory post-synapses in the same cells. **(F)** Zoom on a dendritic segment corresponding to the rectangular area highlighted in (E). **(G)** Confocal image of a primary dendrite from a neuron expressing GFP-actin, further immunolabeled for GFP, showing numerous dendritic spines bulging out of the shaft.

### Modulation of neuroligin-1 expression level affects synapse formation

To assess whether the formation of synapses within neurospheres could be modulated by canonical molecular pathways, we targeted the cell adhesion molecule neuroligin-1 (NLGN1), whose overexpression is known to promote synapse maturation in primary neurons, and whose down-regulation conversely reduces the number of synapses (Chih et al., 2005, 2006; Haas et al., 2018; Levinson et al., 2005; Shipman et al., 2011; Kwon et al., 2012). We thus electroporated neurons with a previously validated shRNA to NLGN1 (Chih et al., 2005; Ducrot et al., 2025; Kwon et al., 2012), or a control shRNA, both containing a GFP reporter. To over-express NLGN1, we electroporated a NLGN1 construct bearing an N-terminal biotin acceptor peptide tag (bAP), plus endoplasmic-reticulum resident biotin ligase (BirA^ER^) inducing its biotinylation (Chamma et al., 2016; Howarth et al., 2005). At DIV14, neurospheres were live stained with streptavidin-AF647 (SA-AF647), then fixed and immunostained with primary antibodies to VGluT1 followed by secondary antibodies conjugated to AF568, and imaged by spinning disk confocal microscopy **(Fig. 7A)**. Although endogenous NLGN1 can localize at both excitatory and inhibitory synapses (Ducrot et al., 2025), we focused here on excitatory synapses because our NLGN1 construct bears both A and B splice sites, thereby expecting to affect mainly the density of VGluT1 puncta (Chih et al., 2006). The number of VGluT1 puncta per unit dendrite area of GFP-expressing neurons was quantified offline **(Fig. 7B)**, and was in the range of 0.255 ± 0.007 per µm² for control neurons (n = 26), in agreement with values previously measured in 2D cultures (Giannone et al., 2013). Excitatory synapse intensity was slightly reduced in neurons expressing shRNA to NLGN1 as compared to neurons expressing the control shRNA **(Fig. 7C,D)**. In contrast, neurons over-expressing NLGN1 formed significantly more excitatory synapses than counterparts expressing the control shRNA alone or the shRNA to NLGN1. When piled together, the spinning disk images of the GFP signals formed a montage of 96 neurospheres **(Fig. S5A)**, illustrating the homogeneous properties and high content capacity of these samples. Interestingly, the maximum intensity projection of the GFP signals showed individual neurons preferentially positioned at the periphery of the neurospheres, while the minimum intensity projection highlighted the more central dendritic network **(Fig. S5B-D)**.

**Figure 7.**
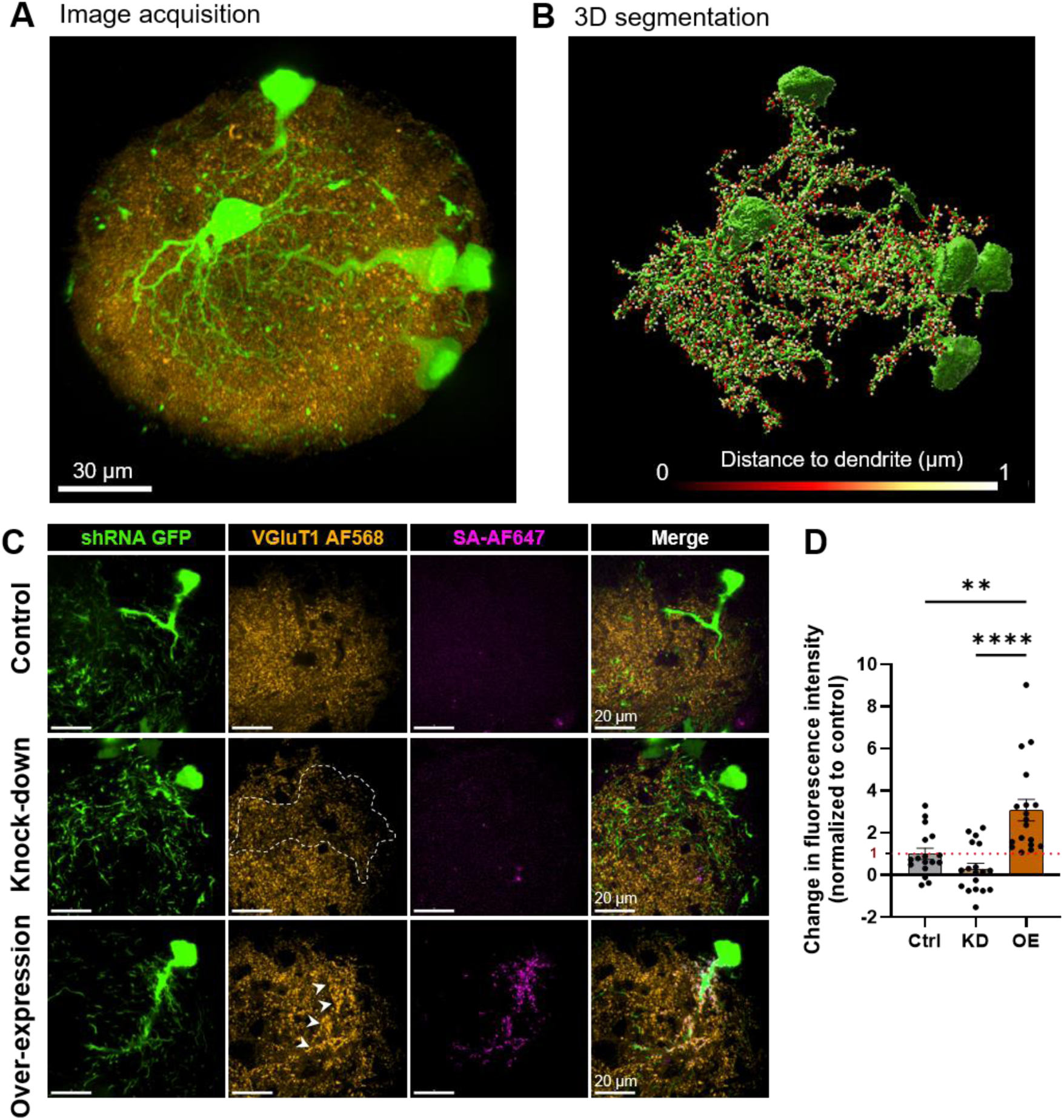
Effect of NLGN1 expression level on the density of excitatory synapses. Dissociated hippocampal neurons were co-electroporated with BirA^ER^ and shRNAs to either a control protein p53 (Ctrl) or to NLGN1, both containing a GFP reporter, together with bAP-NLGN1 in the overexpression condition. **(A)** Maximum intensity projection image of a DIV14 neurosphere, in which some neurons express the GFP reporter (green), while excitatory pre-synapses are immunostained for VGluT1 (orange). **(B)** Corresponding 3D segmentation of the GFP and VGluT1 signals. VGluT1 puncta were selected as belonging to electroporated neurons, based on their shortest distance to GFP dendrites. **(C)** Representative confocal images of shRNA GFP (green), VGluT1 AF568 (orange) and SA-AF647 (magenta) staining in each condition. Note the decrease (dashed line) or increase (arrows) in VGluT1 staining upon NLGN1 knock-down or over-expression, respectively. **(D)** Quantification of the relative change in fluorescence intensity of VGluT1 puncta contacting GFP-positive dendrites compared with control puncta. Values obtained in the KD and OE conditions were normalized to the Ctrl condition, whose mean value was set to 1 (red dotted line). Dots represent individual neurospheres. Data presented as mean ± SEM were compared using a Kruskal–Wallis test followed by Dunn’s multiple-comparison test (n = 15 - 17 neurospheres per condition).

### Imaging of recombinant and endogenous neuroligin-1

We then exploited the rescue condition to examine the surface distribution of recombinant bAP-NLGN1 within dendrites, upon live staining of neurospheres with SA-AF647 **(Fig. 8A,B)**. Only one or two neurons per neurosphere were labelled by SA-AF647, allowing the clear identification of neurites and synapses. Dendrites from individual neurons were decorated with abundant bAP-NLGN1 positive puncta (1613 ± 125 puncta per neuron), with an average synaptic enrichment of 6.5 ± 0.4 (n = 25 neurons). To further improve our resolution on the localization of NLGN1 in the synaptic cleft, we turned to TEM. Moreover, to rule out potential mislocalization artifacts due to the over-expression of recombinant NLGN1, we took advantage of a novel knock-in (KI) mouse strain in which endogenous NLGN1 is N-terminally tagged with bAP that can be enzymatically biotinylated by the exogenous delivery of BirA^ER^, for detection by high-affinity streptavidin conjugates (Ducrot et al., 2025). Neurospheres made with hippocampal cells from KI mice were thus treated at DIV4 with adeno-associated viruses (AAV1s) coding for BirA^ER^ and containing a GFP reporter under an IRES sequence. At DIV14, neurospheres were stained with SA-AF647 for observation in optical microscopy, or to streptavidin conjugated to 1.4 nm gold nanoparticle (SA-Nanogold) for observation in TEM. Neurospheres exhibited strong SA-AF647 signal **(Fig. 8C)**, highlighting the specific staining of biotinylated bAP-NLGN1. Higher resolution observation of neurospheres under confocal microscopy revealed the presence of numerous small puncta, most likely corresponding to bAP-NLGN1 positive synapses **(Fig. 8D)**. In parallel, neurospheres labeled with SA-Nanogold were fixed, silver enhanced, embedded in resin and cut in ultrathin sections before being imaged in TEM. Around 30% of synapses displayed individual nanoparticles, showing the presence of biotinylated endogenous bAP-NLGN1 in the cleft of excitatory synapses **(Fig. 8E)**. We counted 14.1 ± 1.2 nanoparticles per synapse (mean ± SEM, n = 44 synapses), that were more accumulated in the center-half of the synaptic cleft than at the borders **(Fig. 8F)**. bAP-NLGN1 formed a subset of nanodomains (2.0 ± 0.9, mean ± SD, n = 44 synapses), each containing a few nanoparticles, and the number of nanodomains scaled positively with the length of the synaptic cleft **(Fig. 8G)**.

**Figure 8.**
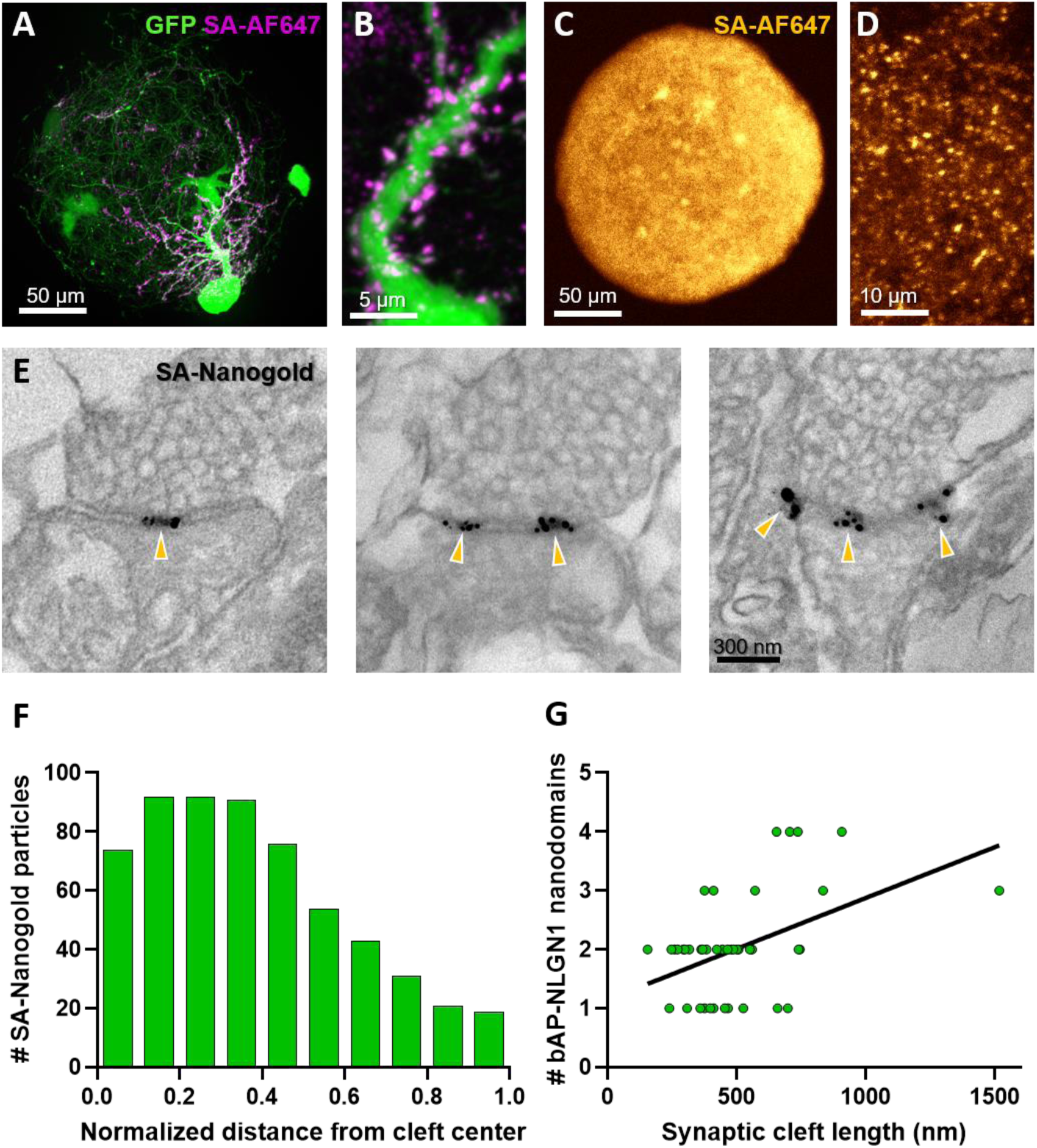
High resolution imaging of recombinant and endogenous NLGN1. **(A)** Maximum intensity projection of a confocal stack taken with a 60x objective, showing an entire neurosphere where neurons have been co-electroporated with shp53-GFP (green), rescue bAP-NLGN1 and BirA^ER^, then live labelled at DIV14 with SA-AF647 (magenta). **(B)** Zoom on a dendritic segment showing numerous synaptic puncta enriched in bAP-NLGN1. **(C)** Epifluorescence image (10x) of a neurosphere formed with hippocampal cells isolated from KI mice, infected with BirA^ER^ AAV1s, and labeled with SA-AF647 at DIV14. **(D)** Higher magnification confocal image (63x), showing many puncta containing biotinylated endogenous bAP-NLGN1 labeled with SA-AF647. **(E)** Representative TEM images of synapses in neurospheres from KI mice infected with BirA^ER^ AAV1s, and labeled with SA-Nanogold before silver enhancement and osmium staining. Examples show 1, 2, or 3 nanodomains of bAP-NLGN1 in the synaptic cleft, respectively (arrowheads). **(F)** Normalized localization of silver enhanced SA-nanogold particles along the synaptic cleft, i.e. zero represents the center of the synapse, and 1 the synaptic border (n = 623 nanoparticles from 44 synapses). **(G)** Occurrence frequency of the number of bAP-NLGN1 nanodomains labeled with SA-Nanogold in the synaptic cleft (each dot represents a single synapse).

## Discussion

In this study, we developed and extensively characterized a 3D culture model made of primary nerve cells isolated from the hippocampi of rat embryos or mouse pups. These cultures are based on the same dissections traditionally done by many laboratories to produce Banker or mixed cultures (Kaech and Banker, 2006). However, when deprived of substrate adhesion in U-bottom ultra-low attachment wells, this mix of neurons and glial cells spontaneously auto-organize into compact spheroids of highly reproducible dimensions and organization. When compared to 2D cultures plated on coverslips, this 3D culture system uses a much smaller number of cells, as well as reduced amounts of tissue culture plastic and media, and is therefore quite cost-effective. Moreover, no evidence of necrosis due to hypoxia or lack of nutrients was found in the center of neurospheres, most likely because of their relatively small size (radius < 100 µm) and porosity, as compared to bigger organoids containing more cells (El Harane et al., 2025). Neurospheres could be observed live, to measure their size, record calcium transients or synaptic currents, but also immunostained in fluid phase to label cellular structures including nuclei, neurites, and synapses. Permeabilized neurospheres were sufficiently accessible to antibodies, and immunolabelled proteins could be detected by confocal microscopy without the need to cut histological sections. However, confocal acquisitions remain fairly long (typically 5-10 min per hemisphere for one wavelength), and one might want in the future to speed up the imaging process, for example using light sheet microscopy (Gao et al., 2019; Beghin et al., 2022). Moreover, since neurospheres readily sediment in resin at the bottom of conical tubes, they represent a nice sample for block trimming and ultrathin sectioning, followed by TEM. Neurospheres are also compatible with the electroporation of DNA constructs prior to seeding (DIV0) or infection by AAVs after spheroid formation (DIV4), allowing for the detection of pre- or post-synaptic proteins. This possibility was exploited to study the effect of the genetic manipulation of NLGN1 on synaptogenesis as well as the synaptic localization of biotinylated endogenous NLGN1.

Using these methodologies, we showed that neurospheres grow in culture, through the combination of glial cell proliferation and the elaboration of a complex network of axons and dendrites, that translates into an increase in the volume of individual cells. The presence of astrocytes in close proximity to neurons, acting through both paracrine release of growth factors and physical contact, is likely to promote neuronal survival and development as well as synapse formation, better than in 2D cultures which remain fragile samples. Moreover, the compliant properties of neurospheres associated to the natural cell-cell interactions that exist in the brain, represent an advantage over neurons grown on rigid substrates coated with artificial adhesive biomolecules such as poly-lysine or laminin. The number of cells, nuclei, and the proportion of neurons versus glial cells could precisely be counted, making neurospheres a highly calibrated cellular system. The projected area of neuronal nuclei was 3-fold larger than that of surrounding astrocytes, as observed in the developing mouse hippocampus (Yazdani et al., Stem Cells 2012). In parallel, growing axons and dendrites eventually formed numerous synapses onto one another. Neurospheres contained both excitatory and inhibitory synapses, as revealed by the presence of the pre-synaptic transporters VGluT1 and VGAT by immunofluorescence, the detection of the post-synaptic scaffolding proteins PSD-95 and gephyrin via electroporated intrabodies, and the recordings of spontaneous EPSCs and IPSCs by electrophysiology, allowing for a balanced excitation to inhibition ratio. We found higher numbers of excitatory synapses versus inhibitory synapses based on both pre- and post-synaptic markers, as previously reported in 2D cultures of dissociated hippocampal neurons (Ducrot et al., 2025).

As a result of sustained glutamatergic activity, neurospheres exhibited periodic calcium oscillations of high amplitude and low frequency (1-2 min^-1^), whose appearance correlated with synapse assembly peaking between DIV7 and DIV10. Fast propagation of the calcium signals throughout individual neurospheres revealed strong local neuronal connectivity. Those calcium pulses might originate from the spontaneous bursts in AMPA receptor-mediated currents detected in electrophysiology, and closely resemble giant depolarizing potentials, a hallmark of hippocampal circuit development in rodents (Cossart and Khazipov, 2022). Individual neurons electroporated with GFP-actin further revealed the presence of multiple well-formed dendritic spines, as observed in organotypic hippocampal slices (Letellier et al., 2018, 2020; Shipman et al., 2011). We also confirmed the synaptogenic function of NLGN1 in neurospheres using a knock-down versus over-expression approach (Kwon et al., 2012; Chih et al., 2005; Levinson et al., 2005) showing a bidirectional modulation of excitatory synapse density. Finally, in live labelling conditions fluorescent streptavidin penetrated neurospheres much better than organotypic slices (Chamma et al., 2016; Ducrot et al., 2025), allowing highly specific detection of biotinylated NLGN1 in synaptic puncta decorating dendrites. We further showed by TEM that endogenous bAP-NLGN1 bound to streptavidin-Nanogold accumulates in the synaptic cleft where it forms a subset of nanodomains, whose number positively scales with the length of the synaptic cleft, as previously demonstrated for HA-NLGN1 and PSD-95 (Nozawa et al., 2022; Cloâtre et al., 2024; Hruska et al., 2018). Assuming that streptavidin saturates biotin binding sites, we estimate that a single excitatory post-synapse contains around 28 bAP-NLGN1 dimers, close to our previous estimate in organotypic slices (Ducrot et al., 2025).

Cellular spheroids have widely been exploited in various domains of life sciences such as cancer research, but not so much in neuroscience yet. Instead, organoids were developed that form layers resembling brain structures (Risbey and Lewandowsky, 2017), but these ‘mini brains’ are tedious to obtain, heterogeneous, require expensive culture media and supplements, and can get contaminated during their long-term culture. In contrast, neurospheres are extremely standardized, low-cost, and rapid to form, making them suitable for drug screening assays to investigate key signalling pathways, for example those governing synapse assembly. Neurospheres can be made from hippocampus, as shown here, but also from cortex (Dingle et al., 2015), and potentially other brain regions that can easily be dissected out (e.g., the striatum), opening the way to the investigation of specific synapse types formed between neurons of different origins, and mixed together. Another interesting extension of this 3D model would be to study the invasion of glioblastoma cells within neurospheres, to further dissect the mechanisms of malignant synapse formation (Venkataramani et al., 2022; Taylor et al., 2023). Finally, a challenge would be to develop 3D cultures of human-derived pluripotent stem cells and differentiate them into neurons (Alessandri et al., 2016; Zhang et al., 2013), allowing the investigation of human-specific proteins, which can have different synaptic functions when compared to rodent counterparts (Marro et al., 2019). We thus hope that this report can stimulate new studies using neurospheres as a model system, that will eventually deepen our understanding of the basic and disease-related mechanisms underlying neuronal connectivity.

## Materials and Methods

### Animals

Pregnant Sprague-Dawley rat females were purchased from Janvier Labs (Saint-Berthevin, France). B6J-Nlgn1^em1(bAP)Ics/Iins^ knock-in (KI) mice, in which the endogenous *Nlgn1* gene was N-terminally flanked by a DNA sequence coding for the 15 aa biotin acceptor peptide (bAP), GLNDIFEAQKIEWHE, were recently described (Ducrot et al., 2025). This mouse strain was generated by PHENOMIN-ICS (Strasbourg, France), then imported and raised to the homozygote phenotype by our animal facility (EOPS). To avoid genetic drift, mice are back-crossed every 10 generations with control mice of the same background (C57/Bl6/J). All procedures involving animals were conducted in accordance with the European guidelines for the care and use of laboratory animals, and the guidelines issued by the University of Bordeaux animal experimental committee (CE50; animal facilities authorizations #A33063940 and ethical project authorizations #21725-2019081114558294). Housing was at a constant temperature (21°C) and humidity (60%), under a fixed 12 h light/dark cycle with food and water available ad libitum.

### DNA plasmids and AAVs

GFP-actin was a gift from A. Matus (FMI, University of Basel) (Fischer et al., 1998). RFP-actin was described previously (Chazeau et al., 2015). The intrabody to PSD-95, Xph20-GFP (Addgene #135,530 pCAG_Xph20-eGFP-CCR5TC), was a gift from M. Sainlos (IINS, University of Bordeaux) (Rimbault et al., 2024). The intrabody to gephyrin, GPHN.FingR-GFP (Addgene # 46296 pCAG_GPHN.FingR-eGFP-CCR5TC), was a gift from D. Arnold (University of Southern California, Los Angeles, CA, USA) (Gross et al., 2013). Short hairpin RNA to murine NLGN1, and its control shRNA to the protein p53 (Chih et al., 2005), were gifts from P. Scheiffele (Biozentrum, Basel, Switzerland). The plasmid coding for BirA^ER^-IRES-GFP was a gift from D. Choquet (Getz et al., 2022; Ducrot et al., 2025). The production of the corresponding AAV1 was made by the viral core facility of the Bordeaux Neurocampus (IMN), obtaining a viral titer of 2.0 x 10^13^ genome-containing particles (GCP)/mL.

### Dissociated rat hippocampal cultures

Hippocampi from E18 rat embryos were dissected out and dissociated into individual cells, as described (Kaech and Banker, 2006; Cloâtre et al., 2024). Cells were counted using an automated fluorescence-based cell counter (LUNA-FLTM, Logos Biosystems). To characterize the impact of the initial number of cells on neurosphere size and nuclei number over culture time, 100’000 non-electroporated neurons were first resuspended in 5 mL warm culture medium made of Neurobasal Plus supplemented with 0.5 mM GlutaMax and B27^TM^ Plus (Gibco #A3582901, #35050038, #A3582801, respectively). Then, 100 µL of this cell suspension was seeded in 24 wells of a U Bottom–Ultra Low Attachment (ULA) 96-well plate (ThermoFisher scientific #174925). Three diluted cell suspensions (1:2; 1:4, and 1:8) were made from this initial stock by sequentially adding 2.5 mL of warm medium into the 2.5 mL cell suspension left over. Thus, the wells contained either 1000, 500, 250, or 125 cells each. For electrophysiology experiments, cells were seeded at densities of 2,000 cells per well in either Neurobasal medium or BrainPhys medium (StemCell Technologies, #05791) supplemented with B27^TM^ Plus. For calcium imaging, neurospheres were made from 500 cells and cultured in BrainPhys medium supplemented with B27^TM^ Plus. For TEM experiments, cells were seeded at a density of 1,000 cells per well and cultured in Neurobasal medium. Neurospheres were kept up to 2 weeks in a 37°C – 5% CO_2_ humidified incubator, and 50 µL of extra culture medium was added to each well after one week.

### Neuron electroporation

To express fluorescent and/or synaptic proteins in neurons prior to immunocytochemistry and confocal microscopy, 300’000 dissociated hippocampal cells were added to 1 mL of warm DMEM + 10 % horse serum (HS) (Eurobio, L0106-500; Gibco, 26050088) and centrifuged at 1,000 RCF for 5 min. The cell pellet was then resuspended in 100 µL of P3 Primary Cell 4D X Kit L solution (Lonza, #V4XP-3024), mixed with 3 µg total DNA (GFP, GFP-actin, Xph20-GFP, GPHN-FingR-GFP; RFP, RFP-actin), transfered in nucleocuvette vessel and electroporated with the Lonza 4D-Nucleofector X Unit using the CU 133 program specific for Neuron rat brain (Hi/Cx High Viability). To modulate NLGN1 expression level, the following plasmid combinations were used: Ctrl: BirA^ER^ + shRNA-P53 GFP (1.5:2 μg DNA); KD: BirA^ER^ + shRNA-NLGN1 GFP (1.5:2 μg DNA); OE: BirA^ER^ + bAP-NLGN1 + shRNA-P53 GFP (1.5:1.5:2 μg DNA). To obtain sparsely labeled cells, neurospheres were formed by mixing electroporated neurons with non-electroporated cells at a 2:3 ratio, and 500 cells of the mix were seeded per U-bottom well in 100 µL culture medium.

### Dissociated hippocampal cultures from bAP-NLGN1 KI mice and infection with AAVs

Hippocampi from B6J-Nlgn1^em1(bAP)Ics/Iins^ KI mice at P0 were dissected out in Hibernate medium and incubated with papain (Sigma-Aldrich, #S3125) for 20 min at 37°C, then mechanically dissociated in Hibernate A (Thermofisher, #A1247501) supplemented with 1.8 mM CaCl_2_. Dissociated neurons were resuspended in Neurobasal A medium (Gibco, #12349-015) supplemented with 0.5 mM GlutaMAX and B27^TM^ Plus, and seeded at a density of 1,000 cells per well of 96 U Bottom Plate -ULA. At DIV3-4, AAV1 coding for ER-resident biotin ligase (BirA^ER^) was added at a concentration of 30,000 MOI per well to each half of a 96-well plate, supplemented with 20 µM D-biotin (Sigma-Aldrich, #B4639). Neurospheres were fed at 7 days with 50 µL per well of Neurobasal A medium and left in culture up to DIV14.

### Measurement of neurosphere size

Neurospheres containing different numbers of initially seeded cells (125, 250, 500, and 1000) were observed live at various time points in culture (DIV 1, 3, 7, and 14) under a shelf microscope (Nikon TS-2) equipped with a 40x/0.55 NA dry objective and a camera driven by the Nikon Imaging System (NIS Element). Images of 5-6 neurospheres in each condition were acquired, and their diameter was quantified offline using the image analysis software Metamorph (Molecular Devices, Sunnyvale, USA).

### Electrophysiology

Neurospheres cultured up to DIV14 in either Neurobasal or BrainPhys media were transferred to a recording chamber, perfused continuously with oxygenated ACSF (in mM: NaCl 126; KCl 2.5; NaH_2_PO_4_.H_2_O 1.25; CaCl_2_.2H_2_O 2; NaHCO_3_ 2; MgSO_4_.7H_2_O 2 and D-Glucose 10) at 28°C, and visualized using an upright video microscope (AxioExaminer Z1; Zeiss) equipped with infrared gradient contrast and a 60X water dipping objective (APO-CHROMAT). Neurospheres were maintained in place using a custom system made of a slice harp on which nylon threads were stretched and attached very close together. Individual neurons were patched using pipettes of resistance 5-8 MΩ prepared from borosilicate glass capillaries (GC150F-10, Clark Electromedical Instuments) with a micropipette puller (P-97; Sutter Instruments, Novato, CA, USA). The patch pipette solution contained (in mM): 135 K-gluconate, 3.8 NaCl, 1 MgCl_2_.6H_2_O, 10 HEPES, 0.1 EGTA, 0.4 Na_2_GTP, 2 Mg1.5ATP, and 5 QX-314 (pH = 7.2, 292 mOsm). The pH and osmolarity of the pipette solution were adjusted at 7.2 and 290 mOsm, respectively. Recordings were made using a Multiclamp 700B amplifier and Digidata 1550B digitizers controlled by Clampex 10.7 (Molecular Devices, Sunnyvale, CA, USA). In whole-cell voltage-clamp recordings, cells were held at −60 mV and the series resistance was monitored by a voltage step of – 5 mV. Signals were low-pass filtered at 2 kHz and sampled at 10 kHz in voltage clamp mode, and at 4 kHz and sampled at 20 kHz in current clamp mode. In voltage-clamp mode, the holding potential was set at −60 mV and successive voltage steps of −5 mV were performed to measure passive properties of the cells including input resistance and capacitance. In current-clamp mode, hyperpolarizing and depolarizing current steps were injected through the patch pipette to assess the active properties of neurons. Current step ranges were −50 to +400 pA, 1000-ms duration. Spontaneous excitatory and inhibitory post-synaptic currents (EPSCs and IPSCs) were recorded in voltage-clamp mode at a fixed membrane potential of −60 mV or +10 mV respectively. To confirm the glutamatergic nature of the EPSCs and the GABAergic nature of the IPSCs, antagonists of AMPA receptors (DNQX, 20 μM, Abcam, Ab120018) or GABA_A_ receptors (gabazine, 1 μM, Tocris Bioscience, SR95531) were respectively used in some experiments. Finally, to confirm the activity-dependence of the synaptic glutamatergic transmission, 1 µM Tetrodotoxin (TTX, Tocris Biosciences #1078/1) was added to the recording bath. Spontaneous events were analyzed using Clampfit 11.1 and a home-made Python script (https://github.com/mlebouc/ABFileAnalyzer).

### Live calcium imaging

A stock solution of Fluo-4 AM calcium indicator (Thermofisher, #F14201) was prepared at 5 mM in DMSO and stored in 2 µL aliquots at -20°C. Neurospheres were loaded with Fluo-4 AM for 10 min at 37°C using a 1:1000 working dilution (5 µM) in culture medium added directly into the wells. Using fetal bovine serum (FBS)-coated pipet tips, neurospheres were collected and rinsed in 10 mL magnesium-free Tyrode solution (15 mM D-glucose, 108 mM NaCl, 5 mM KCl, 2 mM CaCl_2_ and 25 mM HEPES, pH = 7.4) supplemented with 10 µM glycine to potentiate NMDA receptors (Heine et al., 2008), and centrifuged for 3 min at 300 RCF. The supernatant was aspirated and neurospheres were resuspended in 1 mL of the same buffer, then placed in an 18 mm round Ludin chamber. The chamber was gently swirled to concentrate the neurospheres in the middle of the chamber, before being mounted onto an epifluorescence microscope (Nikon Elipse TiE) equipped with a 10X/0.4 NA dry objective, a mercury lamp (Nikon Xcite), and a filter set for Fluo-4 (Excitation: FF01-472/30; Dichroic: FF-495Di02; Emission: FF01-525/30). The microscope is enclosed in a thermostatic box (Live Imaging Services, Basel, Switzerland) that maintains the temperature of the samples at 37°C. Streams of 1,200 images at 100 ms exposure time and binning of 2 were acquired with an sCMOS camera (Prime 95B, Photometrics) driven by the Metamorph software. Spontaneous calcium transients were recorded for several neurospheres at each developmental stage (DIV 3, 7, 10, or 14). To identify the mechanisms underlying calcium dynamics, some neurospheres at DIV14 were exposed to 1 µM TTX (Tocris), 100 µM glutamate, or 40 mM KCl, added 30 s after the start of the recording. Small volumes (20 µL) from concentrated stock solutions were added gently into the chamber to prevent neurospheres from moving. Analysis of calcium signals was performed under the offline Metamorph software. The Fluo-4 intensity averaged over whole neurospheres was corrected for background, normalized by the level at time 0, corrected for photobleaching (< 20%), and plotted over time. The number of calcium peaks above baseline, divided by the recording period was defined as the oscillation frequency (in min^-1^), while the maximal Fluo-4 value was taken as the peak amplitude. For comparison of conditions, (e.g. Control, TTX, glutamate, KCl stimulation), the integrated Fluo-4 intensity above baseline, divided by the recording time, was defined as the calcium influx. Data were collected for each neurosphere and averaged per condition.

### Immunocytochemistry

Neurospheres were collected in a 15 mL Falcon tube, rinsed by adding 10 mL warm Tyrode solution (15 mM D-glucose, 108 mM NaCl, 5 mM KCl, 2 mM MgCl_2_, 2 mM CaCl_2_ and 25 mM HEPES, pH = 7.4, 280 mOsm), and centrifuged at 300 RCF for 3 min in a horizontal swinging bucket centrifuge (Ependorf model 5702). To stain endogenous or over-expressed biotinylated bAP-NLGN1, neurospheres were incubated live with AF647-conjugated streptavidin (Invitrogen, #S32357, 4 μg/mL) diluted 1:500 in Tyrode solution supplemented with 1% biotin-free Bovin Serum Albumin (BSA, Roth, #0163), for 30 min at 37°C, prior to fixation. For all other stainings, neurospheres were gently resuspended in warm PBS containing 4% paraformaldehyde, 4% sucrose and fixed for 2 h at RT. Free PFA binding sites were quenched by 50 mM NH_4_Cl in PBS for 15 min, then neurospheres were permeabilized for 30 min with PBS containing 1% Triton X-100. To block non-specific binding, neurospheres were incubated for 2 h at RT in PBS containing 1% BSA and 0.3% Triton X-100. Neurospheres were incubated with primary antibodies for 24 h at 4°C in a cold room, on an rocking shaker (Starlab, #N2400-8010) with the following references depending on the experiments: chicken anti-GFAP (Abcam, #ab4674, 1:1000) to stain astrocytes, mouse anti-synapsin1 (Synaptic systems, #106 001, 1:1000), guinea pig anti-VGLUT1 (Merck Millipore, #AB5905, 1:2000), or guinea pig anti-VGAT (Synaptic Systems, #131-004, 1:1000) to stain synapses, mixed with rabbit anti-MAP-2 to visualize microtubules (Merck Millipore, #AB5622, 1:800); mouse anti-NeuN (Merck Millipore, #MAB377, 1:200) to stain neuronal cytosol; chicken anti-GFP (Aveslabs, GFP-1020, 1:2500) or guinea pig anti-RFP (Synaptic systems, 390004, 1:600) to amplify GFP- or RFP-fusion proteins, respectively. Neurospheres were then incubated for 24 h in a cold room with the corresponding secondary antibodies, all diluted to 1:1000: goat anti-chicken AF488 (Invitrogen, #A11039), goat anti-mouse AF488 (Invitrogen, #A11001), goat anti-rabbit AF488 (Invitrogen, #A11008), goat anti-guinea pig AF568 (Invitrogen, #A11075), goat anti-mouse AF647 (Invitrogen, #A21236) and goat anti-rabbit AF647 (Invitrogen, #A21244). To stain cell nuclei, neurospheres were incubated for 30 min in PBS containing 1 µg/mL DAPI (Thermofisher, #62248) at RT during the washing step. Washes between incubations were made by filling the tube with PBS or blocking solution (between primary and secondary antibodies) for 10 min followed by centrifugation at 300 RCF for 3 min, while 250 µL of solution was left when aspirating the supernatant to avoid the loss of neurospheres. To prevent neurospheres from sticking during resuspension steps, pipet tips were coated with either pure FBS or anti-Adherence rinsing solution (StemCell Technologies, #01010). Finally, neurospheres were collected and mounted in Fluoromount-G (SouthernBiotech, #0100-01) or RapiClear 1.52 (SUNJin Lab, #RC152001) between a 12 mm coverslip and a microscopy glass slide, using 150 µm thick spacers (SUNJin Lab, #IS015).

### Epifluorescence microscopy

Neurospheres from KI mice where endogenous bAP-NLGN1 was biotinylated with BirA^ER^ AAV1s and further labeled with SA-AF647, were observed under an epifluorescence microscope (Nikon Elipse TiE) equipped with a 10X/0.4 NA dry objective with a mercury lamp (Nikon Xcite) and filter sets for EGFP (Excitation: FF01-472/30; Dichroic: FF-495Di02; Emission: FF01-525/30) and AF647 (Excitation: FF02-628/40; Dichroic: FF-660Di02; Emission: FF01-692/40) (SemROCK). Images of 100-500 ms exposure time were acquired with an sCMOS camera (Prime 95B, Photometrics, Tucson, USA) driven by the Metamorph software.

### Confocal microscopy

Immunolabelled neurospheres were visualized on an upright scanning confocal microscope (Leica SP-5) equipped with 40x/1.40NA or 60x/1.40 NA oil immersion objectives and photomultiplier tubes whose wavelength collecting windows were adjusted to the different labels. Depending on neurosphere size and objective magnification, image formats from 512 x 512 pixels up to 2048 x 2048 pixels were selected. Image stacks were acquired with an accumulation of 2 lines at a 400 Hz scanning speed and a pinhole set to one Airy disk, over half of the neurosphere height using a step size of 1 µm (for nuclei observation), or on sub-regions zoomed on individual dendrites with a step size of 0.3 µm (for synapse observation). Alternatively, for imaging of neurospheres upon modulation of NLGN1 expression level, we used a Leica DMI8 microscope (Leica Microsystems, Wetzlar, Germany) equipped with a spinning disk confocal Scanner Unit CSU-W1 T2 (Yokogawa Electric Corporation, Tokyo, Japan) using an objective HCX PL Apo CS2 63X oil NA 1.4. Images were acquired on a sCMOS Prime 95B camera (Photometrics, Tucson, USA). The LASER diodes used were at 405 nm (100 mW), 488 nm (400 mW), 561 nm (400 mW) and 642 nm (100 mW) for observation of DAPI, GFP, VGluT1 stained with AF568-conjugated antibodies, and bAP-NLGN1 detected with SA-AF647, respectively. Z stacks were done with a galvanometric stage (Leica Microsystems, Wetzlar, Germany). This system was controlled by MetaMorph software.

### Image processing

Quantification of nuclei and synapses in neurospheres was performed using Imaris v10.2.0 (Oxford Instruments). All images were subjected to background subtraction, and a 1.6 gamma correction was applied to the 405-or 488-nm channels to homogenize fluorescence intensity across nuclei or neurons, respectively. Dendrites were segmented using the Surface module, while nuclei and synaptic puncta (VGluT1, VGAT) were segmented using the Spot module with 5-µm or 1-µm reference diameters, respectively. The number of nuclei or synaptic puncta measured in a confocal z-stack of height h was multiplied by the volume ratio between the whole neurosphere (πD_s_^3^/6), where D_s_ is the neurosphere diameter measured from the 2D intensity projection, and that of the truncated neurosphere [πh^2^(D_s_/2-h/3)], to obtain the total number of nuclei or pre-synapses per neurosphere. In experiments where NLGN1 expression was modulated, VGluT1 puncta were assigned to electroporated neurons based on their shortest distance to GFP-positive dendrites using the “Shortest Distance to Surface” function, with a distance threshold set between −1 and 1 µm. Control VGluT1 puncta were selected by defining a region surrounding GFP-positive dendrites and excluding puncta already assigned as contacting these dendrites. For each neurosphere, the mean fluorescence intensity was measured separately for VGluT1 puncta contacting GFP-positive neurons and for control puncta. The percentage increase or decrease in VGluT1 fluorescence intensity at puncta contacting GFP-positive neurons relative to control puncta was then calculated and defined as the change in fluorescence intensity. In experiments where only a few cells were electroporated (Xph20-GFP, GPHN.FingR-GFP, biotinylated bAP-NLGN1 labelled with SA-AF647), the fluorescence signals from maximal 2D projections of confocal stacks were treated with a wavelet-based segmentation program running on Metamorph (Racine et al., 2006), and the total number of detected puncta was divided by the number of electroporated cells.

### Transmission electron microscopy

Neurospheres made of 1000 hippocampal cells from rat embryos were collected in 15 mL tubes, centrifuged at 300 RCF for 3 min, then fixed with 4% PFA-sucrose and 0.2% glutaraldehyde in PBS overnight at 4°C. Neurospheres made of 1000 hippocampal cells from P0 KI mice and infected with BirA^ER^ AAV1s, were collected in separate 15 mL tubes and centrifuged at 300 RCF for 3 min. The neurosphere pellets were resuspended in 12 mL Tyrode solution containing 1% biotin-free Bovine Serum Albumin (BSA, Roth, #0163), and centrifuged again. The final pellets were resuspended in 500 µL Tyrode + 1% BSA and live-labeled for 20 min at 37°C with StreptAvidin conjugated to 1.4 nm NanoGold particles (SA-Nanogold, Nanoprobes Inc., stock 80 µg/mL, 1:100). Finally, neurospheres were rinsed 2 times in Tyrode followed by centrifugation, then fixed with 4% PFA-sucrose and 0.2% glutaraldehyde in PBS overnight at 4°C. The SA-Nanogold was silver enhanced for 10 min at RT in the dark using the LI Silver kit (Nanoprobes Inc. #2012-45ML). All samples were then post-fixed with 1% glutaraldehyde in 0.15 M Sorensen’s phosphate buffer (SPB; Electron Microscopy Sciences #11600-10) for 10 min at RT, then incubated with 1.5% OsO_4_ and 1.5% potassium ferrocyanide for 30 min on ice. Sequential dehydration was performed with 70%, 90%, 95%, and 100% ethanol, followed by one incubation with 100% acetone, then samples were embedded for 2 h with 1:1 acetone and epon resin (Embed-812, Electron Micoscopy Sciences, #14900) in a Beem capsule of conical Tip (Electron Micoscopy Sciences, #69913-05), followed by 100% epon resin at 60°C for 48 h. Trapezoidal blocks of the final neurosphere-rich pellet were trimmed and further cut into 70 nm ultrathin sections with an ultramicrotome (EM UC7, Leica, Germany). Ultrathin sections were then collected on bare 150-mesh copper grids (Delta Microscopy #DG150-Cu) and examined in high contrast mode with a transmission electron microscope (Hitachi H7650, Japan) at 80 kV, equipped with a CCD camera (ORIUS SC1000 11MPx, GATAN Inc., Ametek, USA). Profiles of pre-synaptic boutons were readily identified by the presence of numerous synaptic vesicles. In neurospheres from rat embryos, excitatory synapses were visually identified by the presence of an electron dense post-synaptic density (PSD), while synapses without a PSD were taken as inhibitory. In neurospheres from KI mice stained with SA-Nanogold, the number of silver particles and bAP-NLGN1 nanodomains within each synapse were manually quantified on Image-J by setting a detection threshold for all images, after subtracting the background. The length of the synaptic cleft and the distance of silver enhanced nanoparticles with respect to the center of the synapse were determined manually under Metamorph.

### Statistical analysis

Statistical significance tests were performed using the GraphPad Prism software 10.6.1 (San Diego, CA). Comparisons of data sets with only two conditions were made using a Student t-test if data followed a normal distribution, or a non-parametric Mann-Whitney test otherwise. To compare more than two groups showing normal distributions, a one-way ANOVA followed by Dunnett’s multiple comparisons test was carried out. Alternatively, when criteria for normality were not met, a Kruskal-Wallis test was used, followed by a post-hoc multiple comparison Dunn’s test. Statistical significance is represented as: *p < 0.05, **p < 0.01, ***p < 0.001, ****p < 0.0001. The number of experiments performed and samples examined are indicated in the relevant figure legends. When the differences between groups were not significant, no symbol was indicated on the graphs.

## Acknowledgements

We thank D. Arnold, M. Sainlos, P. Scheiffele, and A. Ting for the gift of DNA plasmids; M. Munier, S. Benquet, and B. Tessier for molecular biology; C. Breillat, S. Daburon, and D. Choquet for viral constructs; A. Béghin for initial experiments; P. Costet, C. Martin, and H. El Oussini at the Plateforme In Vivo Exempte d’Organisme Pathogène Spécifique (PIV-EOPS) of the Bordeaux Neurocampus for maintenance of the mouse strain; D. Gonzales at the genotyping facility of Bordeaux Neurocampus (NeuroCentre Magendie); N. Retailleau, A. Caralp, and L. Leroy at the Cell Biology facility of the Bordeaux Neurocampus (IINS) for the production of rodent primary cultures, N. Dutheil at the Virology platform of the Bordeaux Neurocampus (IMN) for the production of AAVs; M. Petrel, S. Lacomme, and M. Fernandez-Monreal (Bordeaux Imaging Center) for advice on TEM; M. Mondin, J. Teillon, F. Levet, and S. Marais (Bordeaux Imaging Center) for help with confocal image acquisition and/or analysis; A. Castets and Z. Andrieux for logistics (IINS); M Goillandeau (IMN) for the gift of the “Detection mini” software.

This work received funding from Centre National de la Recherche Scientifique (CNRS), Agence Nationale pour la Recherche (grants ANR-21-CE11-0019-01 “Synaptoligation” and ANR-20-CE11-0006-01 “NanoSynAtlas”), Conseil Régional Aquitaine, Fondation pour la Recherche Médicale (Postdoctoral fellowship to C.D. #SPF202209015897), Marie Skłodowska-Curie individual fellowship to C.D. (#101106943), the National infrastructure France BioImaging (grant ANR-24-INBS-0005 FBI BIOGEN), and the GPR BRAIN_2030 from the Bordeaux University Idex program.

## Author contributions

B.C. performed cell electroporation, neurosphere preparation, immunocytochemistry, and calcium imaging; A.D. and T.C. performed cell electroporation, neurosphere preparation, immunocytochemistry, spinning disk acquisition, and image analysis; C.D. carried out viral infection of neurospheres and electron microscopy; L.B. and J.B. carried out patch-clamp recordings and analysis; L.H. performed image analysis; J.B.S. provided software and support; O.T. coordinated the research project, performed epifluorescence and confocal acquisitions, quantified data, and prepared the manuscript draft. All authors discussed the results and participated to the manuscript writing.

## Supplemental Figures

**Figure S1.**
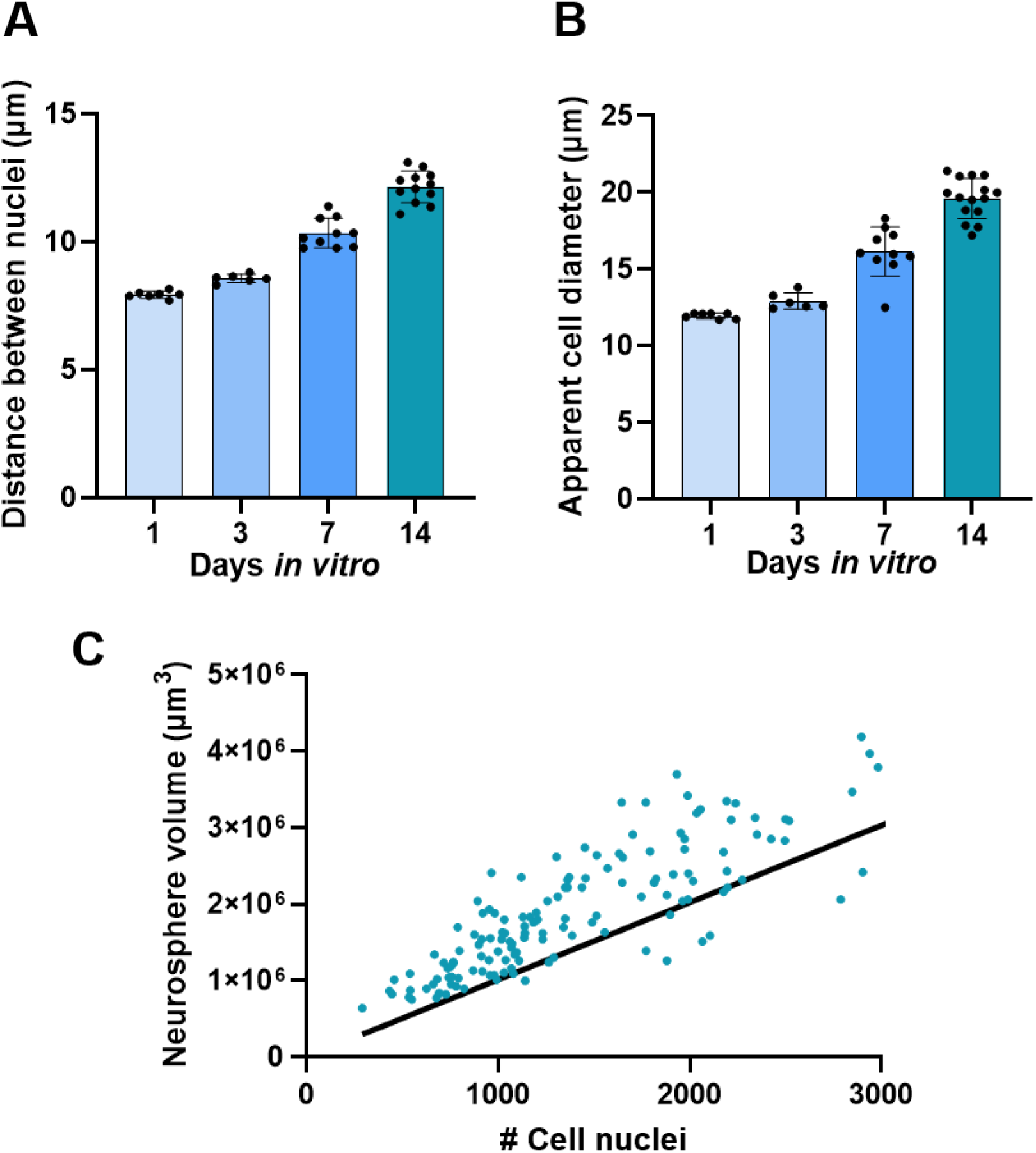
Geometrical parameters of neurospheres. **(A, B)** Distance between the 3 nearest neighbouring nuclei, and apparent cell diameter, as a function of neurosphere culture time, respectively. **(C)** Relationship between the neurosphere volume and the number of cell nuclei it contains. All neurospheres were formed from 500 cells at DIV1, but exhibit some variation in size and cell number when they reach DIV14. The red line is a linear regression through the data (R² = 0.71, P < 0.0001). The mean number of nuclei is 1328 ± 144 (n = 159 neurospheres examined).

**Figure S2.**
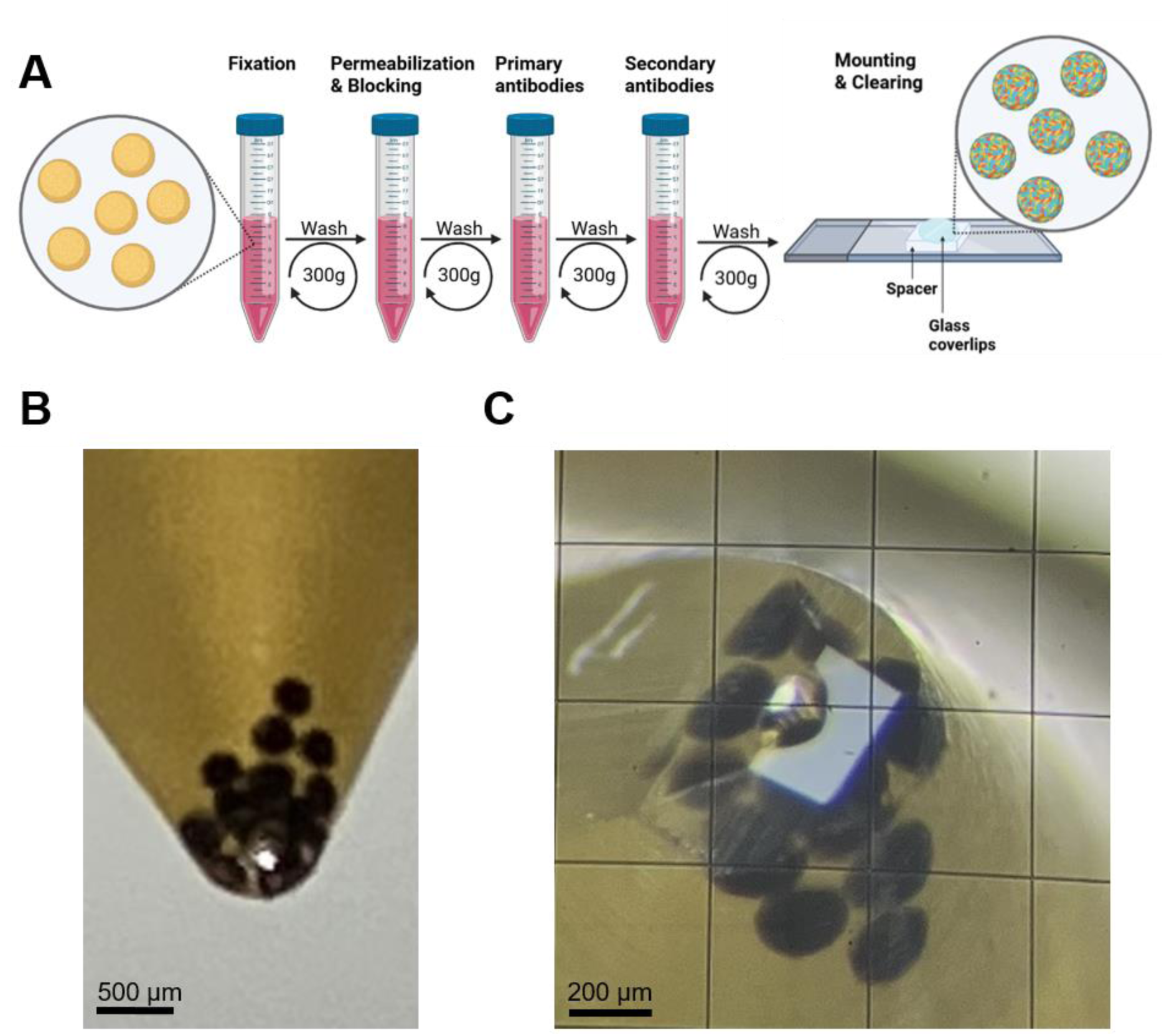
Processing neurospheres for fluorescence and electron microscopy. **(A)** Schematic representation of neurosphere immunolabelling in fluid phase, before mounting the samples for confocal microscopy. **(B)** Image showing a dozen of neurospheres stained with osmium that have sedimented at the bottom of a 500 µL tube filled with resin. **(C)** Illustration of the resin block trimming before ultrathin sectioning prior to TEM.

**Figure S3.**
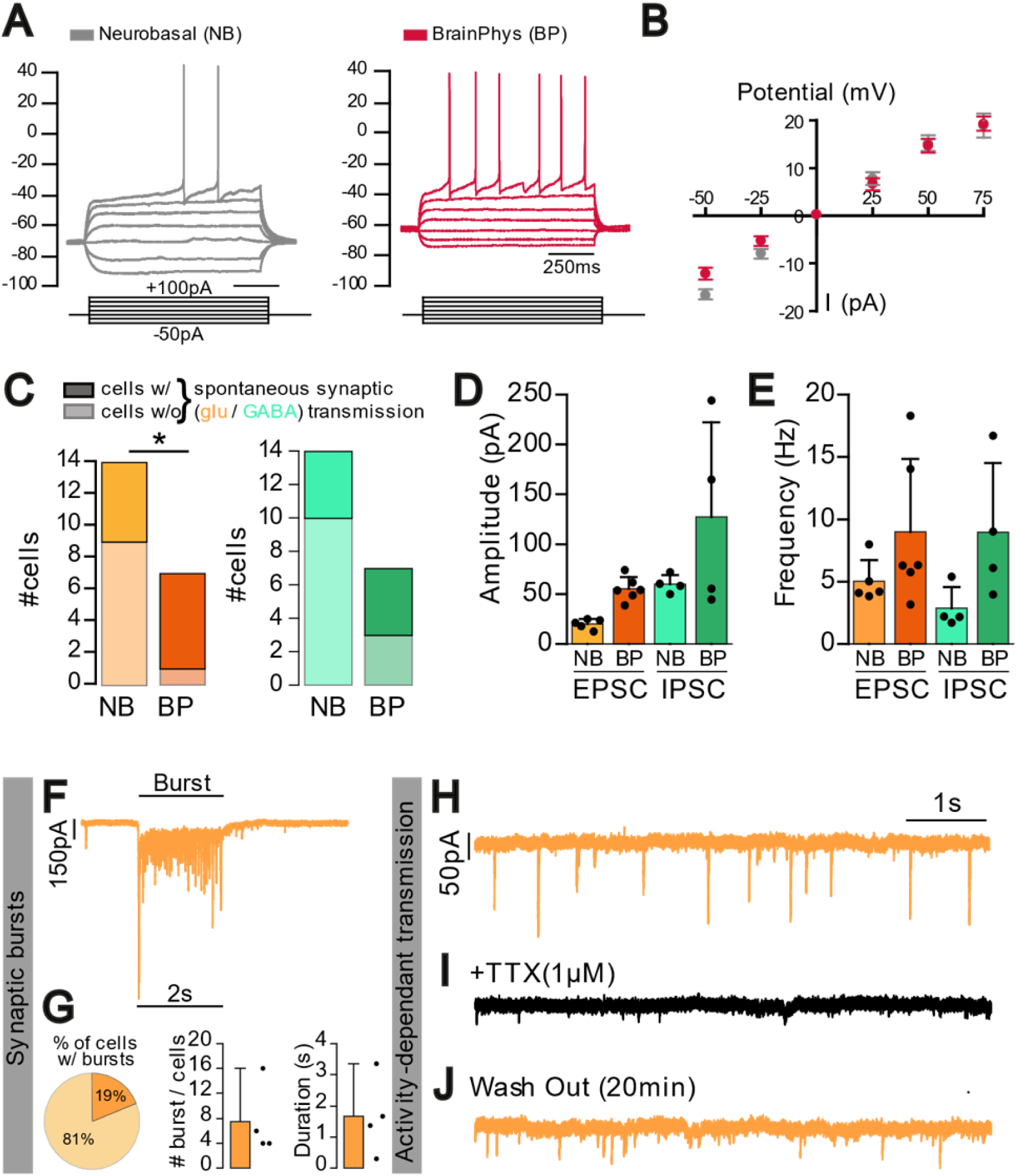
Electrophysiological properties of neurons in Neurobasal versus BrainPhys media. **(A)** Representative traces of membrane potential variations upon incremental current injection (from -50 to 75 pA), for two neurons from neurospheres cultured in Neurobasal or BrainPhys media, respectively. **(B)** Current-voltage (I-V) relationship in Neurobasal medium (grey, n = 11 neurons) versus BrainPhys medium (red, n = 7 neurons). **(C)** Fraction of patched neurons from neurospheres cultured in Neurobasal (NB) or BrainPhys (BP) media, displaying spontaneous EPSCs and IPSCs. Fractions were compared by a Chi² test (* P < 0.05). **(D, E)** Amplitudes and frequency of spontaneous EPSCs and sIPSCs, measured in neurons from neurospheres cultured in Neurobasal or BrainPhys media. **(F)** Example of a burst of glutamatergic synaptic transmission. **(G)** Percentage of cells presenting such bursts among all recorded cells (4/21). Also shown are the number and duration of synaptic bursts per cell in a 200 sec recording period. **(H-J)** Representative traces of activity-dependent glutamatergic synaptic transmission, which is blocked by bath perfusion of TTX and recovered after a 20-min washout. Data are presented as mean ± SEM Dots in population graphs represent individual neurons.

**Figure S4.**
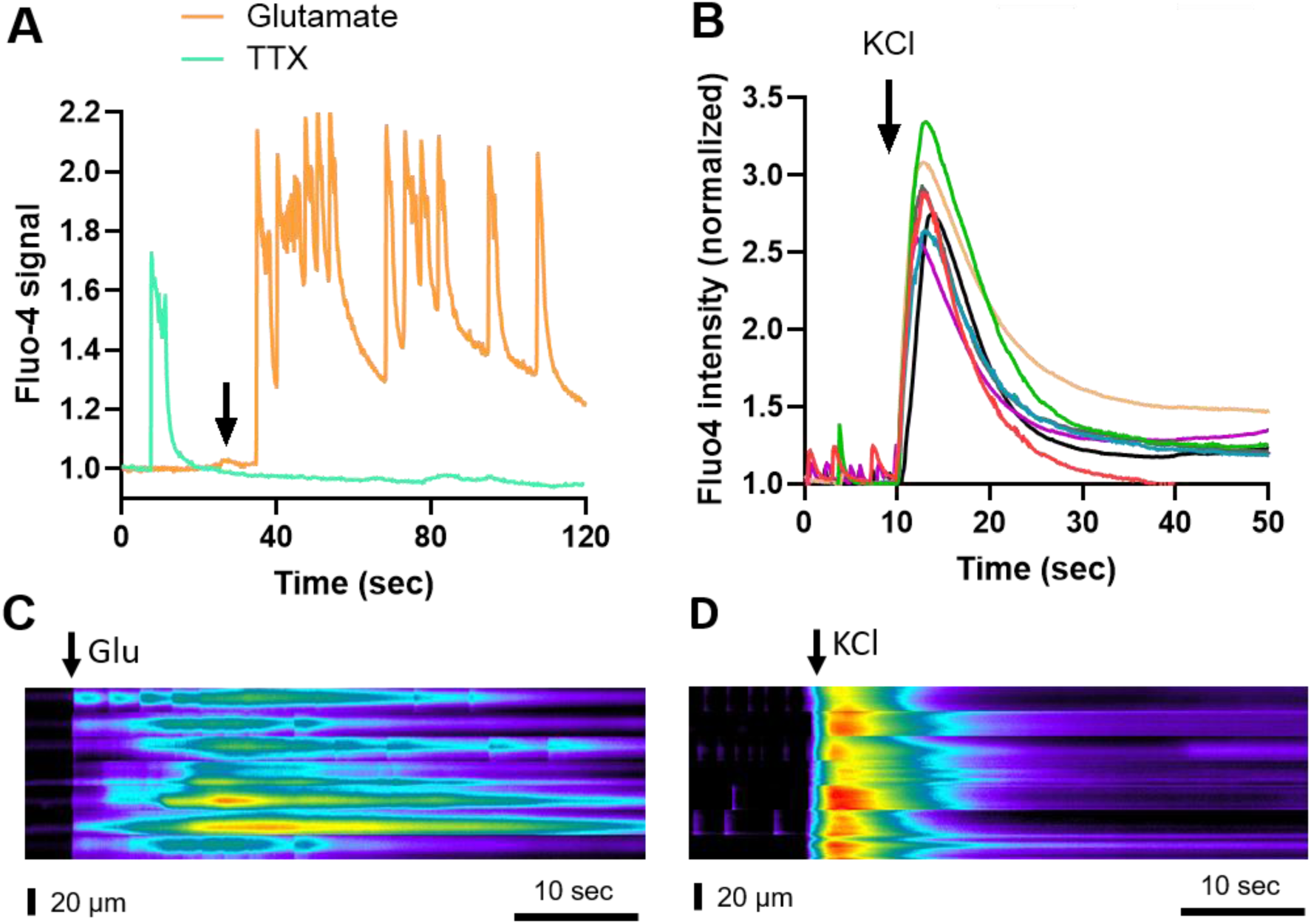
Calcium responses in neurospheres treated with glutamate or KCl. After recordings their spontaneous calcium activity, some neurospheres were exposed to 1 µM TTX, 100 µM glutamate, or 40 mM KCl. **(A)** Fluo-4 signal normalized to the baseline level for two representative neurospheres, upon addition of 1 µM TTX (green) or 100 µM glutamate (orange) at time t = 40 sec (arrow). **(B)** Fluo-4 signal normalized to the baseline level for 6 independent neurospheres, upon addition of 40 mM KCl at time t = 10 sec. All chemicals were left in the bath until the end of the recordings. **(C, D)** Corresponding kymographs. The baseline was set at 0 grey level, and the Fluo-4 intensity in arbitrary units was color-coded. Each line represents an individual neurosphere (n = 8).

**Figure S5.**
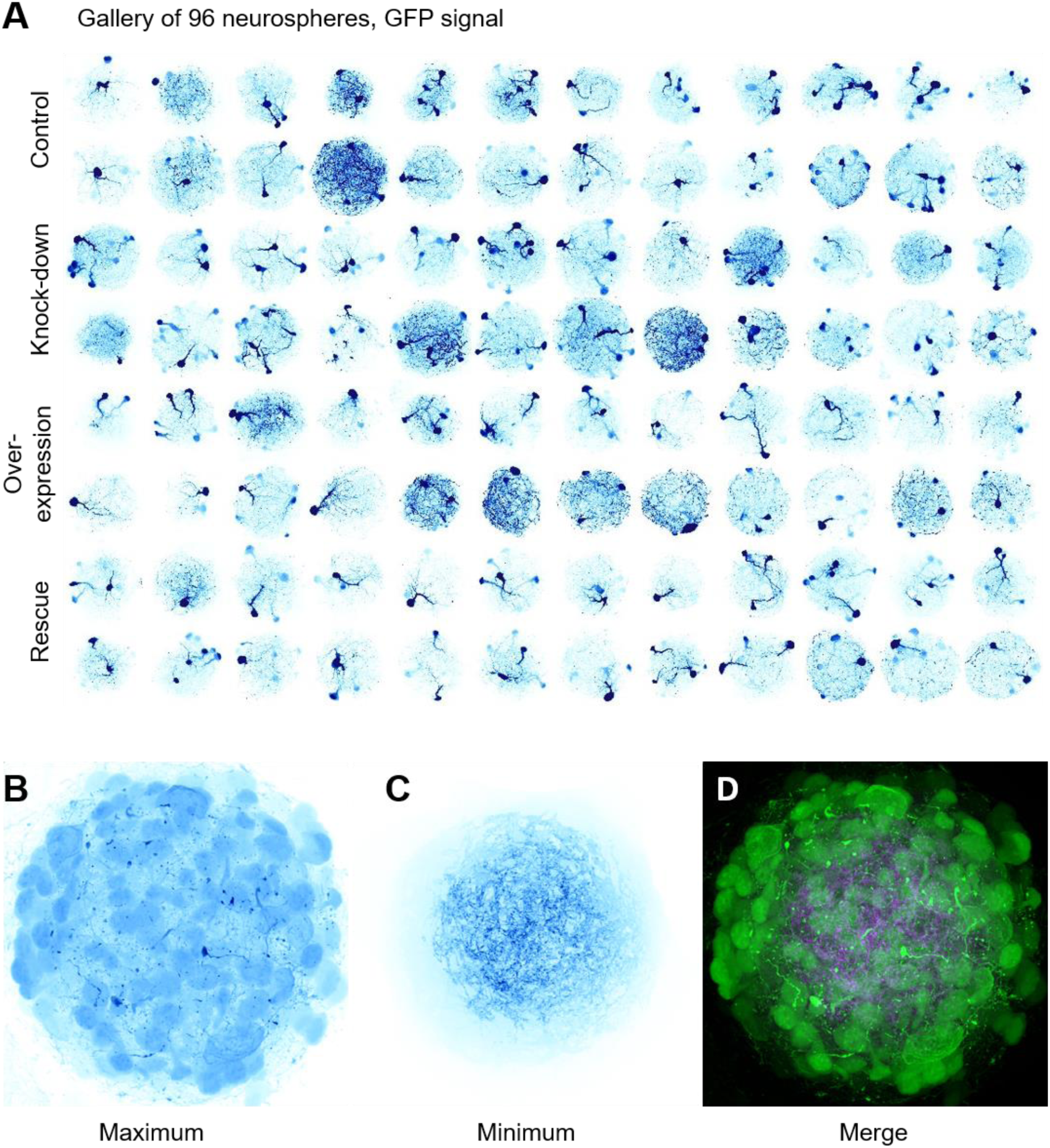
Neurospheres as a high-content culture model. **(A)** Gallery of images showing the GFP signal from 96 independent neurospheres, taken from the different conditions (Control = shp53; knock-down : shNLGN1; Over-expression: shp53 + bAP-NLGN1; Rescue: : shNLGN1 + bAP-NLGN1). Images are color-coded in gold and the LUT is inverted. A gamma correction of 2 was applied. **(B)** Maximal intensity projection of the 96 GFP images showing the cell bodies. **(C)** Minimum intensity projection of the 96 GFP images showing the neurites concentrated in the center of the neurospheres. **(D)** Image merging (C) in green and (D) in magenta.

